# Super-selective reconstruction of causal and direct connectivity with application to *in-vitro* iPSC neuronal networks

**DOI:** 10.1101/2020.04.28.067124

**Authors:** Francesca Puppo, Deborah Pré, Anne Bang, Gabriel A. Silva

## Abstract

Despite advancements in the development of cell-based *in-vitro* neuronal network models, the lack of appropriate computational tools limits their analyses. Methods aimed at deciphering the effective connections between neurons from extracellular spike recordings would increase utility of *in-vitro* local neural circuits, especially for studies of human neural development and disease based on induced pluripotent stem cells (hiPSC). Current techniques allow statistical inference of functional couplings in the network but are fundamentally unable to correctly identify indirect and apparent connections between neurons, generating redundant maps with limited ability to model the causal dynamics of the network. In this paper, we describe a novel mathematically rigorous, model-free method to map effective - direct and causal - connectivity of neuronal networks from multi-electrode array data. The inference algorithm uses a combination of statistical and deterministic indicators which, first, enables identification of all existing functional links in the network and then, reconstructs the directed and causal connection diagram via a super-selective rule enabling highly accurate classification of direct, indirect and apparent links. Our method can be generally applied to the functional characterization of any *in-vitro* neuronal networks. Here, we show that, given its accuracy, it can offer important insights into the functional development of *in-vitro* iPSC-derived neuronal cultures.

## 1. Introduction

*In-vitro* cultures of primary neurons can self-organize into networks that generate spontaneous patterns of activity [10, 74, 58], in some cases resembling aspects of developing brain circuits [37, 24]. The emergent functional states exhibited by these neuronal ensembles have been the focus of attention for many years [15, 83] as they can be used to investigate principles that govern their development and maintenance [35, 44] and to produce biological correlates for neural network modeling [11, 66]. The introduction of human induced-pluripotent stem cell (hiPSC) technologies [69, 82] opened the possibility to generate *in-vitro* neuronal networks in typical [30, 42], as well as patient-specific genetic backgrounds [57, 5, 75, 78, 8, 39], demonstrating the potential to reproduce key molecular and pathophysiological processes in highly controlled, reduced, experimental models that enables the study of neurological disorders and the discovery and testing of drugs, especially in the context of the individual patient [72, 16, 61].

One common approach to obtain information from *in vitro* neuronal networks is to record their activity via multi-electrode array (MEA) or calcium fluorescence imaging and then use network activity features to describe their physiology. One main limitation, however, is that these high-dimensional data, which report about the information representation in the network, do not translate into a clear understanding of how this representation was produced and how it emerged based on neuronal connectivity [14]. The synchronization of spontaneous spike trains among different MEA sites or neurons, also referred to as network bursting, is an example of observed neural behaviors widely reported in the literature. The generation of network bursting in an *in vitro* neuronal culture is evidence that the neurons are synaptically connected. However, the extracellular nature of the MEA recording does not provide information about how neurons are connected and how signals propagate between them, such that computational analyses are necessary to reconstruct their complex dynamic patterns and relate their emergence to the underlying wiring diagram [66]. However, this kind of analysis presents several challenges as it requires not only identification of functional relationships between cells, but also reconstruction of the dynamic causality (i.e., the knowledge of which neuron fires first and affects another one) between directly linked neurons that are simultaneously involved in several different signaling pathways. This defines the difference between functional and effective connectivity inference: the first only reports about statistical dependencies between cells’ activities without giving any information about specific causal and direct effects existing between two neurons [76]; the second attempts to capture a network of effective - direct and causal - effects between neural elements [65].

Only model-based approaches [14, 34] have been proposed for inference of effective connectivity. Among them, dynamic causal modeling (DCM) [18] and structural equation modeling [36] variants have shown best performances. However, these methods estimate the effective connectivity of a measured neuronal network by explicitly modeling the data generation process, i.e. only the connectivity of a simulated network model is inferred without any theoretical guarantee about its accuracy and its ability to correctly estimate the connectivity of the biological network [14, 76].

Because of this limitation, descriptive, model-free approaches are usually preferred as they are easy to implement, rely on a limited number of assumptions that are directly related to the investigated neuronal network, and can be more easily validated [14, 34]. A number of model-free methods proposed for reconstructing the connectivity of *in vitro* neuronal networks [19] have been previously reviewed [48] and tested [76]. However, because they rely on purely statistical indicators, they can only infer how neurons are functionally coupled, but lack the ability to identify the network of effective interactions between neurons by either missing the directionality or confounding indirect and apparent links from direct ones. Directionality conveys the causality of signaling in the network, i.e. which neural element has causal influences over another (Figure 1A(i)). However, causality does not imply a direct connection between two neurons. In fact, a functional coupling between two neurons can be causal even though the two neurons are not directly connected, and this may occur if there is a multi-neurons pathway between the two cells (indirect connection, Figure 1A(ii)), or if the connection detected between the two neurons is simply a mathematical artifact resulting from the correlation of correlations generated by common inputs from other participating neurons (apparent connection, Figure 1A(iii)) [17].

**Figure 1:**
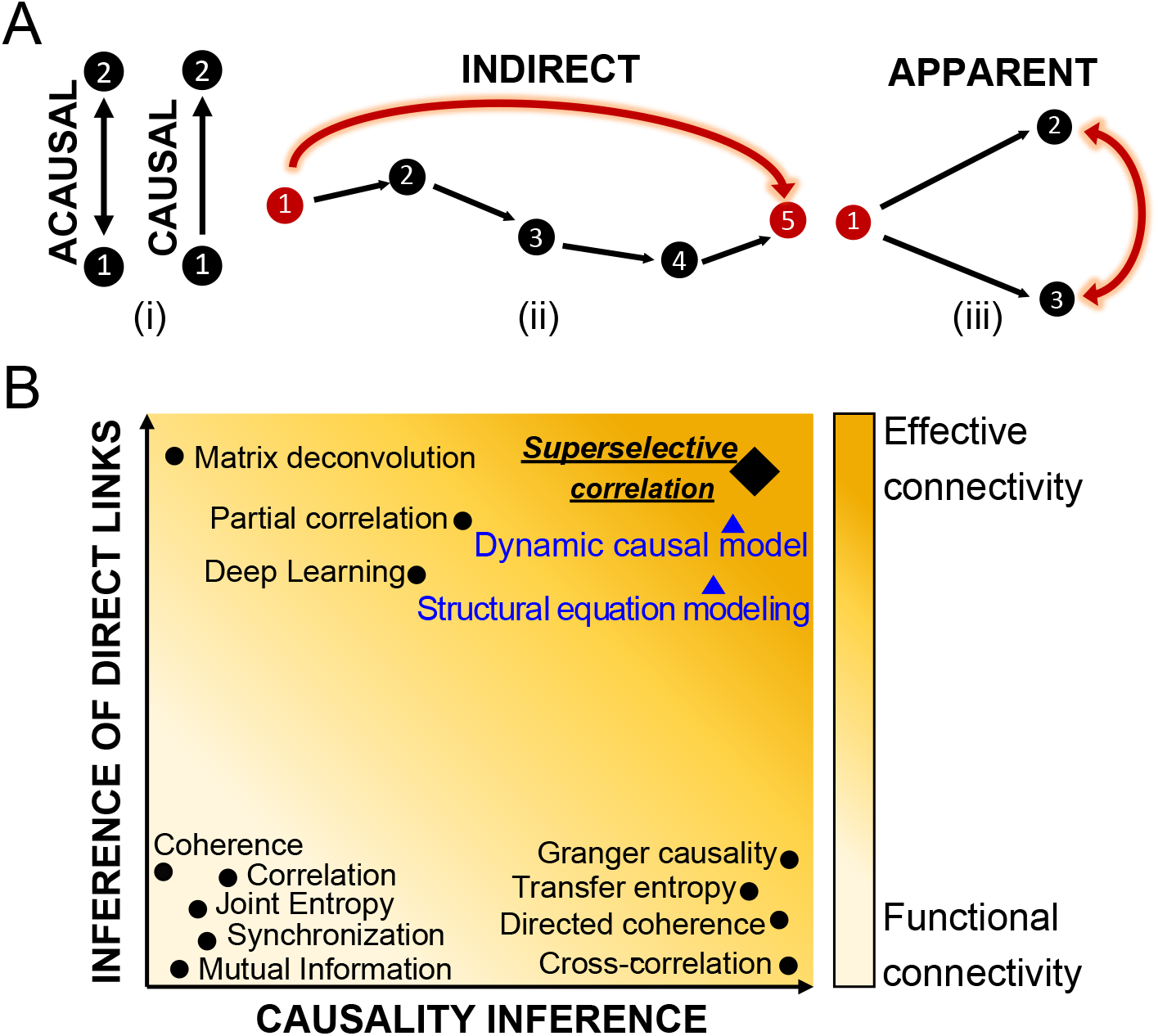
Definition of functional and effective connectivity. (A) Classification of causal (directional link), indirect (multi-neurons pathway) and apparent (functional coupling due to common input) connectivity. (B) Classification of most common connectivity inference methods in terms of causality and detection of direct links. On the *x*-axes, the graph shows a scale of causality which refers to the ability of a given connectivity method to infer or not the directionality of the functional connections between neurons. On the *y*-axes, the graph visually quantifies the capabilities of one approach to detect direct links between neurons by identifying and discarding multi-neurons connections and apparent ones. Indicators that are acausal and do not infer direct links can only report about functional connectivity (light yellow); indicators which contain information about direction of interaction and direct neuron-to-neuron communication are close to the inference of effective connectivity (orange color). Most common model-free techniques are indicated with black dots. Model-based methods are reported with blue triangles and their ability to infer effective connectivity is negatively weighted by the impossibility to test their performances. The super-selective correlation approach we propose is reported in black like the other model-free methods; a diamond signal is used to emphasize the fact that, although being model-free, it aims at inferring effective connections. The graph visually summarizes results of comparisons between connectivity methods from review papers [14, 76, 19] and do not contain precise quantitative information about the differences.

Methods such as correlation [53], coherence [25, 23, 12], mutual information [19, 22, 51], phase and generalized synchronization [51, 2], and joint entropy [19] describe only statistical dependencies between recorded neurons without carrying any information of causality or discriminating direct and indirect effects. Techniques such as cross-correlation [19, 28], directed and partial directed coherence [1, 56], transfer entropy [19, 28, 33] and Granger causality [21, 59] are examples of causal indicators as they provide inference of directionality of dependence between time series based on time or phase shifts, or prediction measures. However, because these operators rely only on pairwise statistical comparisons and treats pairs of neurons independently, they show the same limitations when dealing with indirect connections and external inputs. Only a few techniques can compete in the challenge of inferring the effective connectivity of a network. Partial-correlation [19, 68], which takes into account all neurons in the network, showed best performance in detecting direct associations between neurons and filtering out spurious ones [45]. The most significant limitation of this solution is its high computational cost. Moreover, as the partial correlation matrix is symmetric, this method is not useful for detecting the causal direction of neuronal links. It also does not attempt to infer self-connections [14]. A combination of correlation and network deconvolution was used by Magrans and Nowe [13] to infer a network of undirected connections with elimination of arbitrary path lengths caused by indirect effects. However, this method also can not identify directions of connections and the singular value decomposition of network deconvolution has an extremely high computational complexity [45]. A convolutional neural network approach [54] showed the same limitations in computational complexity and undetected self- and causal connections. Figure 1B graphically summarizes the inference capabilities of the state-of-the-art connectivity methods as reported in [14, 76, 17, 2].

In this work, we propose a novel, mathematically rigorous method that uses a model-free approach (i.e. does not depend on a set of underlying assumptions about the biology of participating cells) to decompose the complex neural activity of a network into a set of numerically validated direct, causal dependencies between the active component neurons that make up the network. First, the inference power of statistical approaches (signal-, network- and information theory- based) allows mapping the functional connectivity of the network. Then, we propose a mathematically rigorous selection scheme that distinguishes between apparent or non-direct links and direct ones, therefore enabling inference of direct causal relationships between connected neurons which more realistically describe the effective connectivity of the network (Figure 2).

**Figure 2:**
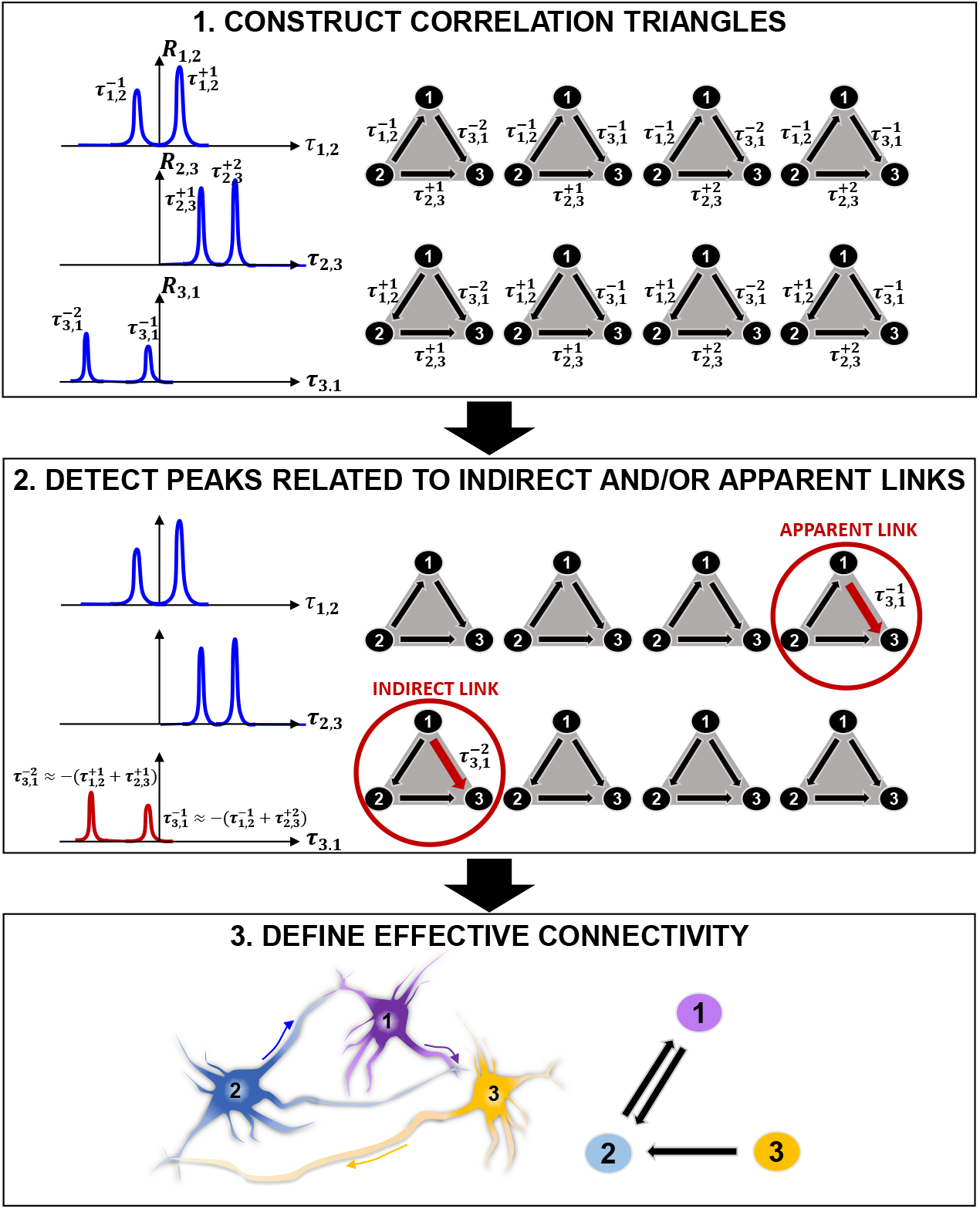
Connectivity reconstruction via detection of correlation triangles and classification of indirect and apparent links. . 1. Given three neurons 1, 2 and 3, our algorithm searches, in time, for all correlations between them by computing the pairwise correlation functions *R*_1,2_, *R*_2,3_ and *R*_3,1_. In this representative example, the algorithm detects two correlation peaks for each pair of neurons and associates the corresponding delays of interactions 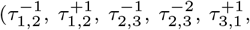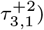 which are defined by the location of the peaks with respect to the origin of the *x* axis. 8 possible combinations of interactions can occur in time between the three pairs of neurons. These correspond to the 8 correlation triangles shown on the right side of the panel. 2. Among the correlation triangles, the algorithm detects 2 critical cases 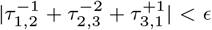 and 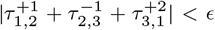 and identifies the peaks relative to an indirect (multi-neurons path) and an apparent (common output) connection by searching for the ones with smaller amplitude. The smallest peaks (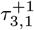 and 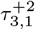) are discarded from the analysis. 3. The correlation triangles are functional to the estimation of the direct connections in the network. Because no more correlation exists between neuron **1**and **3**, the estimated effective connectivity includes only the direct links for (1, 2) and (3, 1): two connections with opposite directionality exist between neuron 1 and 2 because positive and negative correlation peaks are detected in *R*_1,2_ (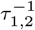 and 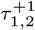); one link connects neuron 2 to neuron 3 as a result of the positive correlation peaks in *R*_2,3_ (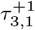 and 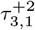).

We evaluate the performances of the proposed method on synthetic datasets generated through simulation of an integrate-and-fire neuronal network mimicking the activity of *in vitro* cultures of neurons, and demonstrate important improvements, relative to the state-of-the-art connectivity methods, to network inference accuracy due to a deterministic component of our method capable of identifying false positive connections.

We show an experimental application of our approach to spontaneously generated *in vitro* networks of human iPSC-derived neurons cultured on MEAs providing an analysis and interpretation of the physiology not possible otherwise. We describe the temporal evolution associated with the connectivity and dynamic signaling of developing hiPSC-derived neuronal networks, including increasing synchronized activity and the formation of small numbers of hyper-connected hub-like nodes, as similarly reported by others [30, 8]. These results further support the performance quality of our approach and provide an example of how this connectivity method can be used to characterize network formation and dynamics, thus facilitating efforts to generate predictive models for neurological disease, drug discovery and neural network modeling.

## 2. Materials and Methods

### Theoretical framework for connectivity reconstruction

The central contribution of this manuscript is in providing an innovative, computationally efficient, and easy-to-apply method for decomposing the collective firing properties stored in the electrophysiological recordings from neuronal networks on MEAs into direct (one-to-one) and causal (directional) relationships between all participating neurons. We propose a multi-phase approach that identifies and discards any correlation link that does not directly relate to a direct interaction between two cells. The core of our methodology is graphically described in Figure 2 and includes three main phases: 1. statistical, correlation-based reconstruction of functional connectivity; 2. mathematically-rigorous super-selection of direct links via identification of peaks related to indirect and apparent links and 3. reconstruction of directed causal connectivity between neurons.

1. The functional connectivity (statistical dependencies) of a network is computed via pairwise correlation studies. Functional interactions between neurons are represented by correlation peaks and their delays ***τ***. The algorithm constructs ***correlation triangles*** by considering all possible combinations of correlation delays for any possible triplet of neurons (Figure 2.1). Importantly, correlation triangles do not refer to any three-neuron physical connection, sometimes referred to as “neural triangles” in the literature [63, 55]. Here we define a correlation triangle as a mathematical object that our algorithm uses to classify functional interactions based on all possible triplets of correlation delays that can be formed in the network. Therefore, correlation triangles exploit the entire signal history of neurons in order to determine the correlation peaks.
2. Correlation peaks associated with indirect or apparent links in corresponding correlation triangles are discarded from the analysis by means of a mathematical super-selection rule which deterministically classifies the type of dependence between each triplets of neurons (Figure 2.2). The super-selection rule is formally presented later. Here, it can be summarized as follows. If a correlation triangle is made up of three correlation delays that are the combination of one another, one of the component correlation delays is either representative of an indirect link (Figure 1A(ii)) or of an apparent link (Figure 1A(iii)); therefore, this correlation delay does not refer to an effective connection and must be discarded. When the algorithm finds a correlation triangle which satisfies this condition, it deepens into the classification of the involved correlation delays and selects the correlation peak to remove based on the peak’s amplitude. For example, in Figure 2.2, the algorithm identifies an indirect link between 1 and 3 (a multi-neuron pathway), and an apparent link between 1 and 3 (correlation due to a common output). The correlation peaks corresponding to these links in the correlation triangles are discarded from the analysis. Importantly, the algorithm removes correlations from the analysis, but does not remove inferred physical connections.
3. Only when all correlation peaks between two neurons are discarded, the algorithm recognizes that a specific interaction is only apparent and deletes the corresponding connection. For example, in Figure 2.3, there is no existing connection between neurons 1 and 3. The estimated effective connectivity includes only direct links for (1, 2) and (2, 3): two connections with opposite directionality exist between neuron 1 and 2 because positive and negative correlation peaks are detected in *R*_1,2_ (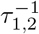 and 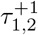); one link connects neuron 2 to neuron 3 as a result of the positive correlation peaks in *R*_2,3_ (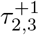 and 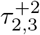).

**Figure 3:**
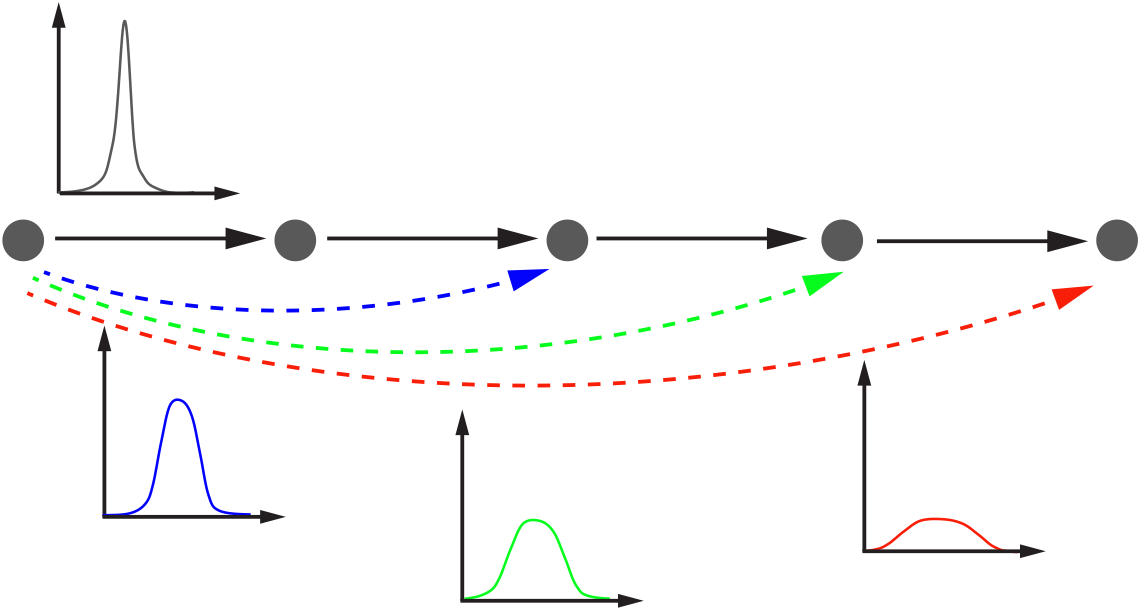
The effect of indirect connectivity on correlation peaks. The edge AB, BC, CD, and DE are directed. The associated correlation peaks are high and narrow because there is a direct dependency between the activity of neuron A(B,C,D or E) and the one of neuron B(C,D or E). The links AC, AD and AE are indirect, with increasing order of correlation between the neurons because of more intermediary cells on the paths that link A to C, D and E. The longer the path, the weaker the dependency between the neurons which results in a lower and wider correlation peak.

The following sections describe the mathematical details of the developed technique. The connectivity reconstruction algorithm and associated functions for performance evaluation were implemented in Matlab and code is available online at https://github.com/fpuppo/ECRtools.git.

### Reconstruction of functional connectivity

#### Temporal correlations

To identify the temporal correlations between the activity of all pairs of *N* recorded neurons *j, k* ∈ {1,…,*N*} in the network, we computed the pairwise correlation function between the corresponding signals *s_j_* and *s_k_*

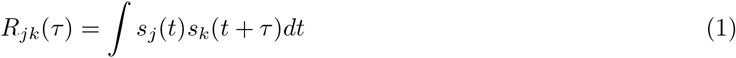

In this formulation, the indexes *j* and *k* are restricted to *k* > *j* in order to avoid unnecessary calculation of auto-correlations (*j* = *k*) and explicitly calculate correlations only for *k* > *j* because, thanks to the symmetry of (1),

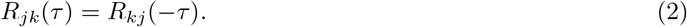

Using the Fast Fourier Transform (FFT) and the correlation theorem, computing correlations in Eq. (1) can be efficiently performed in *O*(*S* log(*S*)) with *S* the number of samples composing the signals *s_j_*. In fact, if 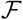 is the FFT operator and 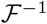 the corresponding inverse, the correlations can be efficiently computed as 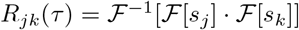 [4].

Peaks of *R_jk_*(*τ*) represent correlations among neurons *j* and *k*. Their amplitude can be regarded as a measure of the level of correlation between the spikes in their registered firing activities. The higher the amplitude of a peak in *R_jk_*(*τ*) (if any), the higher the probability that there is a statistical dependency between neuron *j* and neuron *k*. However, as explained earlier in the text, the existence of a functional coupling between two neurons does not necessarily imply that there is an effective connection between them. For example, the activity of any of the two neurons can have an effect on the other through one or more interconnecting cells between them (Figure 1A(ii)).

The location of each correlation peak with respect to the origin indicates the temporal delay 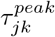 between the activity of neuron *j* and neuron *k* (Figure S1). The sign of this delay, i.e. whether the peak is found on the positive or the negative quadrant of the correlation function, defines the directionality of the interaction that, in the ideal case of a direct connection, suggests which is the pre- and post-synaptic neuron in the interaction.

#### Peak detection

We implemented a peak detection algorithm applied to *R_jk_*(*τ*) to identify all existing functional correlations between any pair of neurons (*j, k*) in the network and to discern the directional dependency between their spiking activities.

As part of the peak detection phase, we used a smoothing Gaussian filtering [62] applied directly to *R_jk_*(*τ*) to remove high frequencies and facilitate the proper identification of correlation peaks.

As introduced above, we assume that a correlation peak in *R_jk_*(*τ*) represents a potential connection between *j* and *k* and that 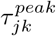 is the signal propagation delay between them. We define a temporal range (−*T*, +*T*) over which to perform the peak search.

For a given parameterization of the Gaussian filter and a defined time window (−*T,* +*T*), the peak detection algorithm allows us to identify a list of pairwise temporal delays 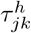 between all pairs of neurons (*j, k*) in the observed network, with *h* ∈ {−*h_n_,…,* −1, 1,…, *h_p_*}_*jk*_ where *h_n_* is the number of peaks with 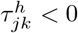 and *h_p_* the number of peaks with 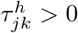. Peaks identified on the positive ({1,…*h_p_*}_*jk*_) or negative ({−*h_n_*,…, −1}_*jk*_) side indicate whether the spiking activity of neuron *j* has temporally occurred, respectively, before 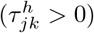 or or after 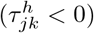 the firing of neuron *k*. Finally, from Eq. (2) the following relation holds:

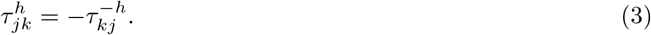

### Detection of false positive connections

Correlation peaks detected in *R_jk_*(*τ*) represent any type of statistical dependence between two neurons. Peaks relative to functional dependencies due to multi-neuron connections or apparent coupling are the main cause of false positives generated in the connectivity reconstruction process and a major source of error in competing methods. To address this, we introduce a framework that identifies the effective network configuration via implementation of a deterministic super-selection rule over all the detected correlation triangles.

### Pure direct connections and correlation triangles

In order to identify possible dependencies between correlations, i.e., if a correlation among neurons is not direct but results from a third party correlation, we consider *cyclic* triplets of correlation delays 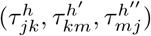. Here cyclic means that each neuron’s index appears in two ordered neuron’s index pairs, once as the first index and once as the second index. This cyclicity directly implies that, if we had

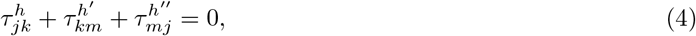

one of the three delays would result from signals that correlate through an intermediary signal as shown in Fig. 2.

However, since the correlations among the neurons’ firing are not Dirac deltas but rather Gaussian-like, we can define a threshold *ϵ* > 0 such that a weak version of (4) still holds. Therefore Eq. (4) reads

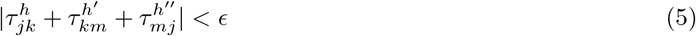

This equation identifies the case where there is a near-perfect match between the correlation delay of each of the three cells and represents the first step in the proposed super-selection algorithm. The definition of “near-perfect match” is based on the choice for the parameter *ϵ* which represents the acceptable degree of temporal approximation when computing Eq. 5. In ideal conditions, *ϵ* → 0 and Eq.(5) reduces to Eq.(4).

Finally, it is useful to define a correlation triangle as the triplet 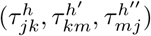 of cyclic correlation delays that the algorithm detects for any pair of neurons in the network. The algorithm aims to construct *n_jk_* × *n_km_* × *n_mj_* correlations triangles given by all possible combinations of correlation peaks of any triplet of neurons’ pairs (Figure 2).

To have a complete picture of our method, we can define the directed graph *G* = (*V, E*) whose vertexes *V* are the active neurons and edges *E* are the active synaptic connections. This definition implies that our algorithm reconstructs only the connections between the neurons in the culture whose activity has been recorded, and contains no reference to the full structural connectivity of the biological network under study. Within this framework, each correlation triangle shares up to three edges with *G*, and the union *E*′ of all 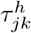 defines the direct graph *G*′ = (*V, E*′). It follows that

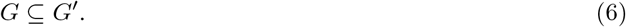

*G*′ can overestimate *G* because it could include connections that are false positives (Figure 2A). For this reason, we need to determine an efficient minimization scheme to reduce *G*′ as close as possible to *G*.

### Edge covering minimization

In order to reduce *G*′ to *G* it is sufficient that for each correlation triangle satisfying Eq. (4) (or Eq. (5)) we identify the dependent 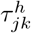 and remove it from *E*′. Therefore, the challenge is to find a discrimination rule that allows us to select the correct 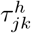 in the correlation triangles that satisfy Eq. (4) (or Eq. (5)).

By considering the nature of neuronal signals, we can define the discrimination factor based on the amplitude 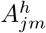 of correlation peaks. In fact, a neuron is equivalent to an input-output object that generates an output signal *s_out_*(*t*) either independently or dependently on an input signal *s_in_*(*t*) coming from another neuron. If *s_out_* depends on *s_in_*, the two signals are not perfectly synchronized but are often noisy (have a phase noise) resulting from the intrinsic excitability properties of the neurons. For example, when an incoming spike train from an input neuron activates an output neuron, the timing of the output spiking depends on many biochemical factors including for example the state of voltage gated ion channels. The result is that the signal of the output neuron is never triggered at the same exact delayed time, but varies. The larger the variation of the delayed timing between the input and the output neurons, the larger the phase noise in the associated correlation peak which will have smaller amplitude and larger standard deviation than the correlation among neurons with input and output signals without phase noise and perfectly synchronized. Moreover, if the signals belong to two neurons that are interconnected via intermediary cells, the phase noise is amplified and the correlation peak is even shorter (low amplitude) and wider (large standard deviation) (Figure 3).

Our method takes into consideration only triangles because a similar analysis performed on higher degree polygons would be redundant. To clarify this point, let’s consider the example reported in Figure 4. Let’s take a graph composed of four nodes. If we consider the polygon ABCD, then there will be a dependency between the edges (the correlation delays) iff

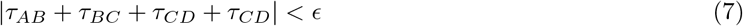

**Figure 4:**
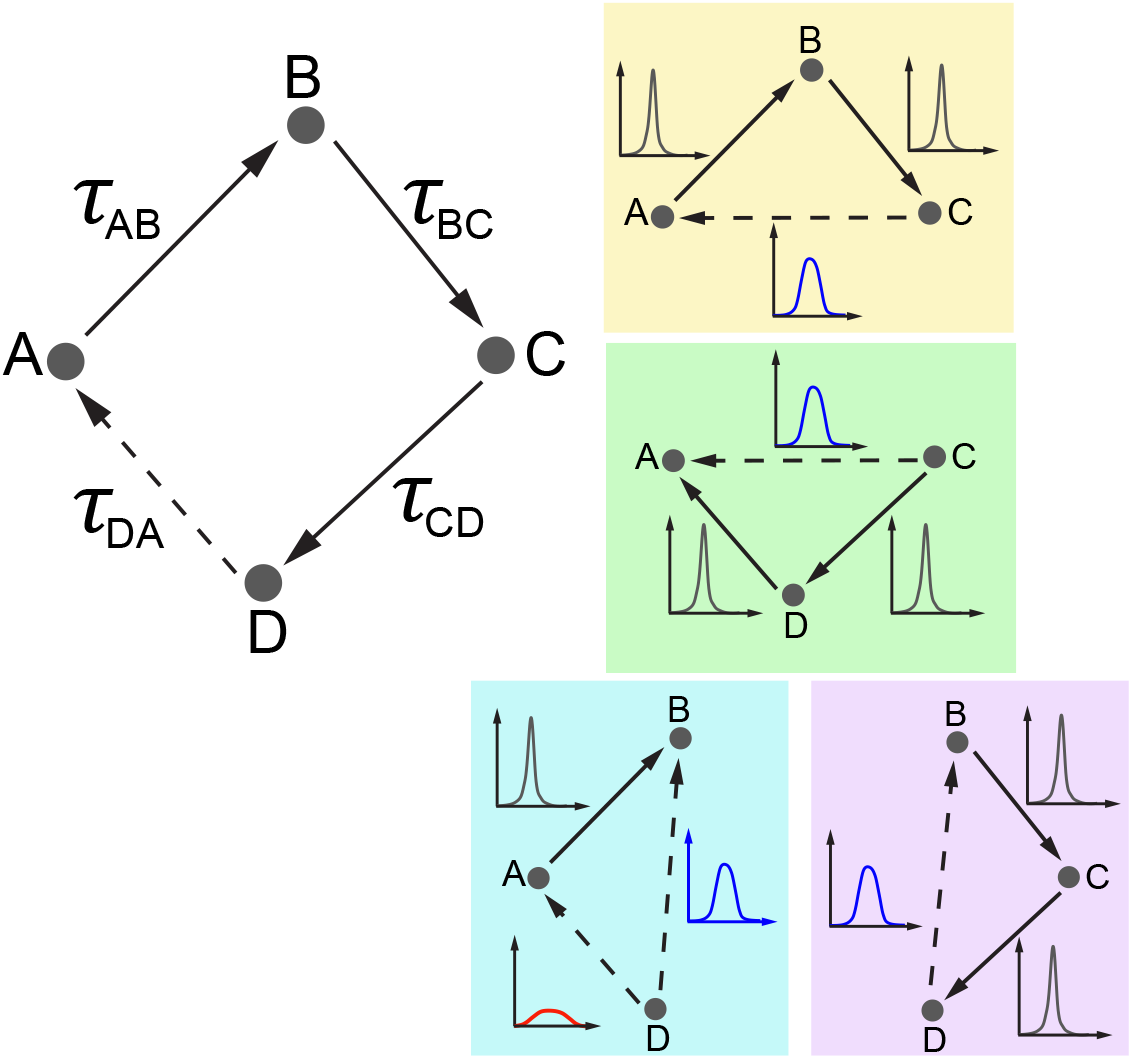
Correlation triangles reconstruct all possible dependencies between neurons. On the left, a graph composed of four neurons A, B, C and D (polygon ABCD). Each edge is associated with a correlation peak and corresponding correlation delay *τ_AB_*, *τ_BC_*, *τ_CD_*, and *τ_DA_*. True connections are indicated by solid lines. The delay *τ_DA_* corresponds to an indirect connection which the algorithm must identify and discard. This is indicated by a dotted line. On the right, the polygon ABCD is decomposed in four triangles ABC, ACD, ABD, and BCD (each color represents a different triplet). The participating correlation triangles reconstruct all possible dependencies among all neurons in the polygon ABCD. Each correlation link is associated with a correlation peak whose amplitude and width depends on the length of the indirect correlation path connecting the two correlated neurons (see Figure 3). In triangles ACD and ABD, the delay *τ_DA_* results from a signal that has propagated through three links AB, BC and CD. Thus, *τ_DA_* will be detected and discarded because having lower amplitude and wider peak than the other peaks.

Therefore, one of these edges is dependent and the algorithm should discard it from the analysis. Let’s say that, for example, *τ_DA_* is the indirect connection. If we consider the triangles ACD or ABD as in Figure 4, in both cases, the delay *τ_DA_* is the one detected and discarded because, as explained in Figure 3, the corresponding correlation peak has lower amplitude and larger width as it results from a signal that has propagated through three links.

Within this picture, we can establish the second step in the super-selection algorithm as:

*Given a correlation triangle 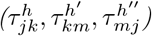, the delay associated with the smallest correlation peak amplitude*

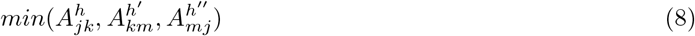

*estimates the false positive connection and is discarded from E*′.

In the ideal case, i.e. without errors and approximations, this super-selection scheme eliminates all indirect correlations, and therefore reduces *G*′ exactly to *G*.

Figure 5 reports a real case scenario of apparent connection. Three neurons (indexed 1, 4 and 12 in our model) from a recorded biological neuronal network are temporally related as described by the corresponding pairwise correlations (Figure 5A) and the resulting correlation triangle schema visualized in Figure 5B. The algorithm checks all correlation triangles in the recorded network and detects that this particular case satisfies Eq. 5 for a selected *ϵ* =3 ms. As explained in the following section, *E* was taken equal to the mean width of the correlation peaks. One correlation delay matches the combination of the other two. The algorithm identifies which of the three peaks corresponds to an indirect or apparent link by comparing the peaks’ amplitude. In the example, the delay *τ*_1,4_ corresponds to the smallest correlation peak and is therefore discarded from the analysis, i.e., from *E*′. If in the chosen temporal window that defines *E*′ only *τ*_1,4_ is detected for neuron 1 and 4, the final reconstruction will not include any effective connection between 1 and 4. This is a nice example of marrying-parents effect [14] where an apparent link between neuron 1 and 4 is formed as a result of neuron 12 firing at the same time on both of them.

**Figure 5:**
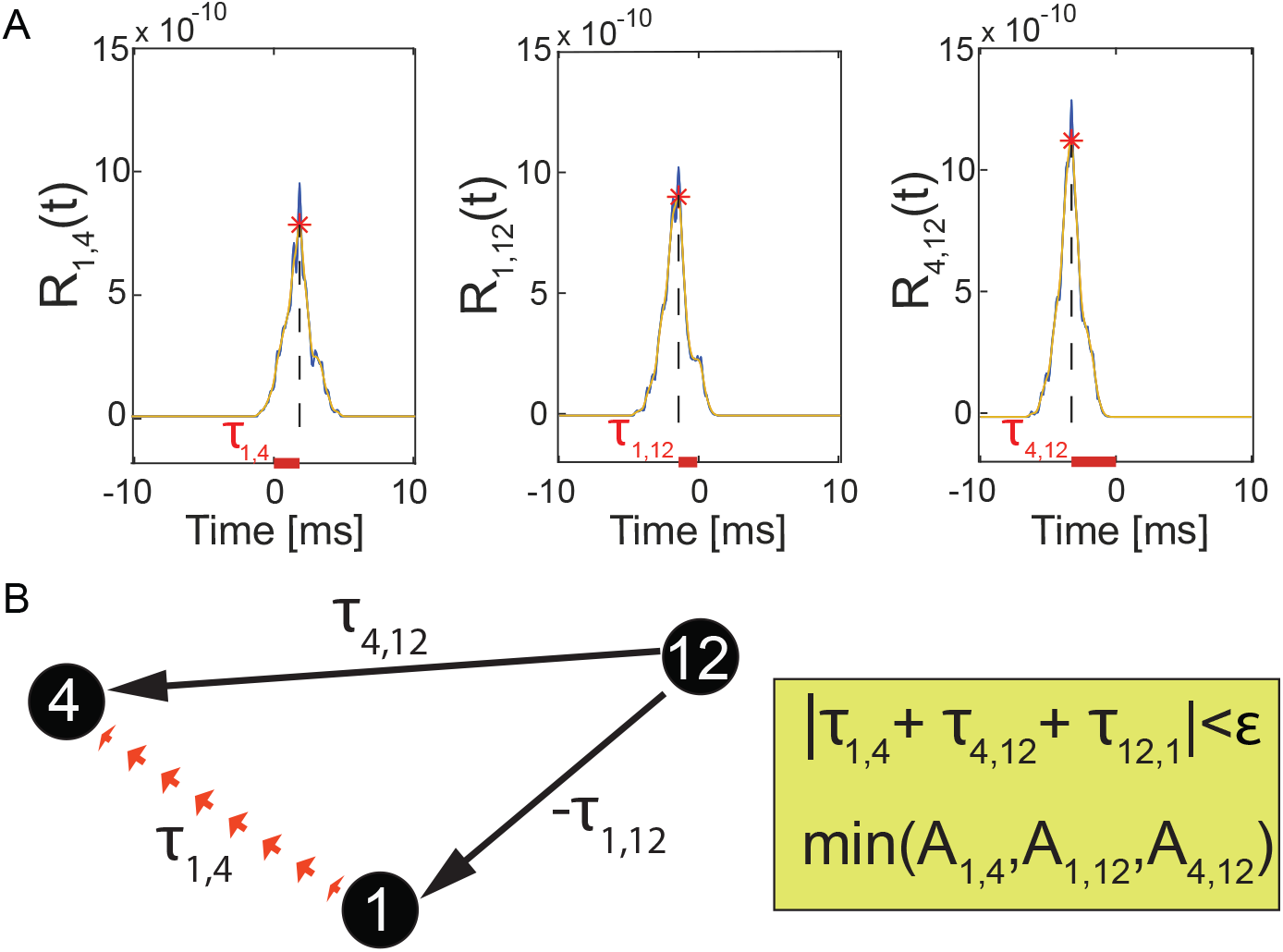
Real case scenario of correlation triangles and detection of false positive connections. (A) Correlation functions *R*_1,4_, *R*_1,12_, *R*_4,12_ computed for pairs of biological neurons (1, 4), (1, 12), (4, 12) (blue line) connected as in panel B. The selected time window was (−20 ms, +20 ms), but the graph only shows a zoomed view in (−10 ms, +10 ms) to better describe the selection rule if only one correlation was detected in time between the neurons. The *σ* of the Gaussian filter was 0.0005 s, resulting in smoothed correlation signals (orange dashed line). The detected correlation peaks (red stars) represent functional correlations between neurons and their combinations define the correlation triangles that our algorithm uses to detect false positive connections. The horizontal offset between the position of each peak and the origin represents the temporal delay *τ_j,k_* associated with these links. (C) These delays satisfy the cyclic condition on *τ*, i.e., |*τ*_1,4_ +*τ*_4,12_ +*τ*_12,1_| = |*τ*_1,4_ +*τ*_4,12_ − *τ*_1,12_| < 3ms therefore indicating that one of the three correlations in the correlation triangle corresponds to an apparent link. The algorithm detects which of the three by searching for the correlation peak with smallest amplitude. In the example, *A*_1,4_ has the smallest amplitude and the algorithm discards it from the analysis preventing generation of a false positive link.

### Connectivity matrix reconstruction

There are three fundamental parameters that affect the inference performance of this method:

- *T* defines the time window (−*T*, +*T*) over which to search for correlation peaks in the correlation function *R_jk_*. This parameter affects the filtering power of the connectivity algorithm. An optimal *T* depends on the specific activity properties of the neuronal network under analysis. To account for all direct neural interactions, the best time window should include the mean maximum propagation delay between the neurons in the network. If the window is too small, the algorithm can over-filter otherwise important correlation interactions.
- *σ* is the standard deviation (width) of the smoothing Gaussian filter and defines the frequencies to filter out in the correlation functions. *σ* is important because the location of the detected correlation peaks weakly depends on it. In fact, while the Gaussian filter is necessary for a more reliable peak detection, the level of smoothing introduces a small temporal jitter between the actual location of the peak and its filtered version.
- *ϵ* is the degree of uncertainty; it defines the threshold beyond which the correlation delays in a correlation triangle can be considered the combination of one another, i.e. how similar the combination of two correlation delays *τ_km_* and *τ_mj_* must be from the direct correlation delay *τ_jk_* to estimate a three-neuron effective connectivity (see Eq. 5). Large *ϵ* values increase the number of detected correlation peaks and computed correlation triangles with the potential shortcoming that true positive connections are filtered out. Too small *ϵ* values are responsible for a poor filtering of spurious delays. A good approximation for *ϵ* is the mean width of the detected correlation peaks which represents the variance of the correlation delays. For example, if we consider the correlation peaks from the real case scenario in Figure 5, the width of the peaks is about 2.5 ms; therefore, we chose *ϵ* = 3 ms as threshold value for our super-selection of direct links.

All three parameters should be tuned in order to have the best outcome from our method.

The connectivity reconstruction approach we propose here is based on a parameter variation scheme that leads directly to the reconstruction of the effective connectivity matrix of the network. We used *TP/N_c_* (net number of true positive (TP) connections, with *N_c_* the number of connections in the simulated neuronal network), *FP/N_c_* (net number of false positive (FP) connections) and Δ = (*TP* − *FP*)*/N_c_* (confidence indicator) to investigate the effect of *T*, *σ* and *E* (Figure 6A-C). The evaluation of the minimized *G*′*p* enabled identification of 80-95% of the total positive direct connections present in the model (*TP/N_c_*) in a wide range of *p* = (*σ, T*), as demonstrated by the curve plateau in Figure 6A. The remaining percentage of connections corresponded to the net number of false connections (*FP/N_c_*) that the algorithm was unable to sort out (Figure 6B) which remains very small for most parameter values. For small *T* (*T* < 2 ms), we observed a decay in sensitivity clearly due to over-filtering of true direct correlations. When doing so, the false positives first increased due to complete failure of the algorithm in recognizing connections in the small temporal range of observation and then rapidly decayed for very small *T* because no peaks could be found. This demonstrates the importance of choosing larger values both for *T* and for *σ* rather than smaller ones in order to avoid missing any correlation information that could negatively bias the algorithm performances. Figure 6C shows the resulting variation of Δ which reached a peak of confidence at 65% and was maintained constant for a wide range of parameters.

**Figure 6:**
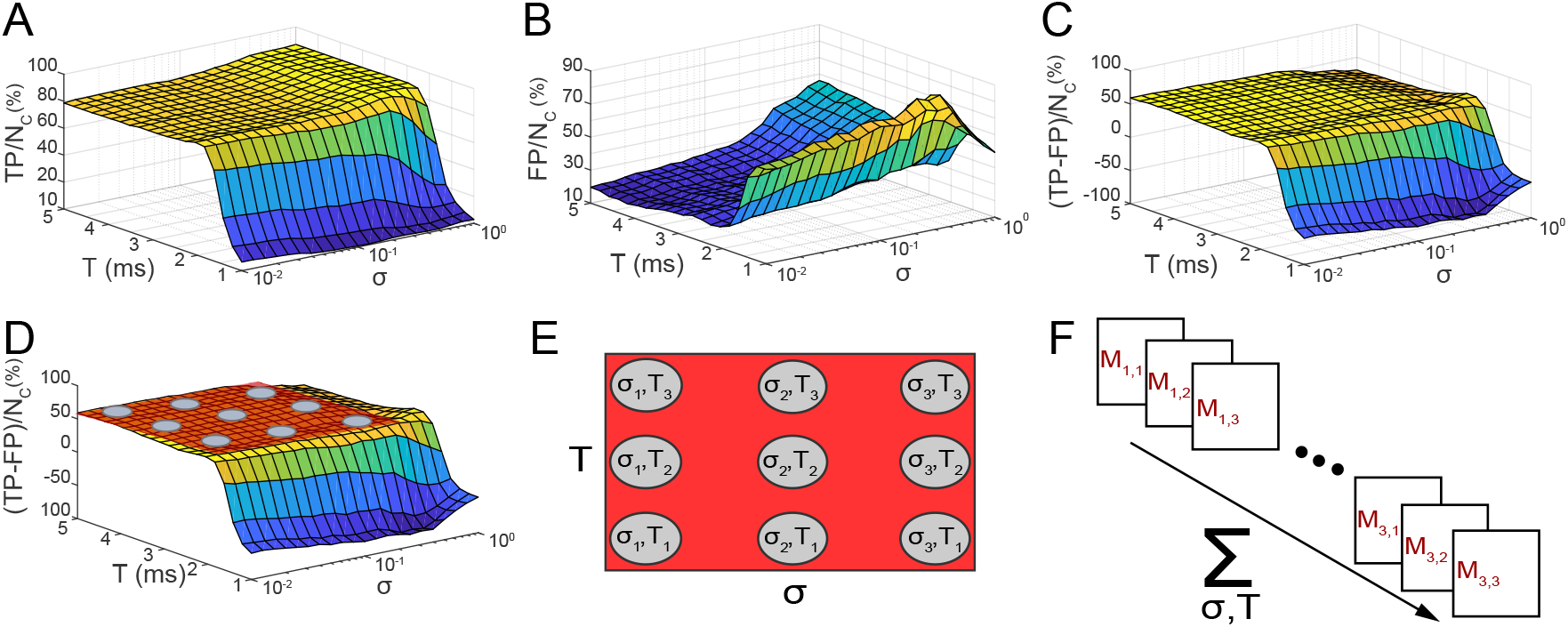
Connectivity reconstruction. (A-C) Evaluation of the performance of the connectivity method under varying time window (−*T,* +*T*), standard deviation *σ*, and for a fixed = 0.7 ms, for one simulated network of 10 neurons. (A) Net number of detected true positives *T P/N_c_*. (B) Net number of false positives *FP/Nc*. (C) Ratio Δ = (*T P* − *FP*)*/N_c_* as chosen metric for evaluation. *N_c_* is the known total number of connections in the simulated network. (D) Visual highlight on the behavior of Δ: a peak of performance is reached around 65%; low variability is proven for a wide range of *T* and *σ* (plateau indicated by the red plane). (E) Definition of a collection of *K* = 9 points *p* in the *T*-*σ* space for which the algorithm shows performances falling in correspondence of the plateau area. (F) Abstract representation of the statistical method for recognition of false positives connections and connectivity reconstruction. For each point a connectivity matrix *Mp* is computed, resulting in the computation of *K* = 9 different connectivity matrices for the same input network. Combination of the results enables computation of the frequency *f_jk_* for each connection following Eq. 9.

This process allowed us to individuate a range of values for *T* and *σ* where the inference performances of the algorithm reached a maximum plateau (Fig. 6D), and define the reconstruction rule of our connectivity method as follows (the numerical details are discussed in the next section of Materials and Methods).

For a fixed value of *E*, we consider a collection of *p* points in the *T* - *σ* space (for example the nine points in Fig. 6D,E). The boundaries for *σ* and *T* can be chosen according to their definitions. For example, a minimum value for *σ* should be related to the very high frequency in the signal while the maximum value to the frequencies in the lower middle spectrum. A minimum value of *T* should be at least as large as 2-3 times the average delay among neurons to guarantee that the relevant peak are included in the analysis. On the other hand, the maximum value for *T* can be chosen as several times (for example 5-6) the average delay among neurons in order to include more correlation triangles later used to refine the selection.

Then, for each point *p* = (*T, σ*) we compute 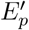 and perform the super-selection to minimize 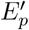. Then, we compute the connectivity matrix *M_p_* (e.g. Fig. 6F) with 1 in the (*j, k*) entry if the reduced 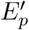 contains at least one 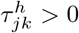, and 0 otherwise. We consider only 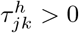 because, according to Eq. (3), if 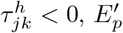 includes 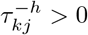 that corresponds to the same correlations.

We can therefore define the frequency for each connection in the connectivity matrix as

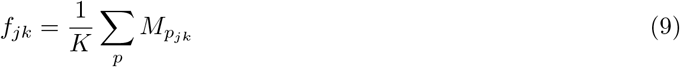

where *K* is the total number of computed points *p*. Each frequency computed in this way is a binary classifier for a given effective connection between neurons *j* and *k*. Therefore, introducing a discrimination threshold 0 ≤ *d* ≤ 1, for each connection we can decide if it is true or false and compute the fraction of true and false positive connections.

### Experimental methodology

#### Synthetic neuronal network model

To develop and validate our connectivity method we used spiking data generated via simulations of neuronal networks based on the Izhikevich model [29] (see Supplementary Material and Fig. S2). The original code was modified to guarantee high levels of activity in the network as well as bursting like behavior similar to that registered in our experiments (Figure S2). We performed our analysis on sparse networks (*N_c_ « N_tot_*, with *N_c_* the number of connections in the simulated network and *N_tot_* = *TP* + *TN* + *FP* + *FN* is the total number of possible connections that can be formed given the input size of the investigated network model) that could be more easily simulated and analyzed with standard computational resources and that accurately described the sparse activity of the iPSC-derived neuronal networks we investigated. However, it is worth noting that our model is general and not restricted to sparse connectivity.

To evaluate the scaling properties of our method, we used data generated via simulations of network models having a varying size of 10, 20, and 50 nodes. For each network size, 20 different networks were randomly generated, simulated and then analyzed for connectivity reconstruction. For each tested network we compared the adjacency matrix reconstructed for a frequency *f_jk_* to the input connectivity matrix of the simulated model.

### Performance measures

The performances were assessed based on the indicators *TP/N_c_* (net number of true positive (TP) connections, with *N_c_* the number of connections in the simulated neuronal network), *FP/N_c_* (net number of false positive (FP) connections) and Δ = (*TP* − *FP*)*/N_c_* (confidence indicator). Δ is independent on the connectivity of the network being reconstructed and, because of its definition, it better defines the level of confidence in the detection of TPs by highlighting the method capabilities in rejecting or not the false positive connections. For the sake of comparison with the literature, we also used the more standard Accuracy measure *ACC* = (*TP* + *FP*)*/N_tot_* [49, 19].

### Numerical experiments

We used *TP/N_c_*, *FP/N_c_* and Δ to investigate the effect of *T*, *σ* and *ϵ* (Figure 6A-C): from pre-selected ranges of these parameters based on observations on the simulated activity, we tuned *T*, *σ* and *ϵ* by looking at the method’s performance in selected ranges (*T* ∈ (1,5) ms, *ϵ* = 0.7 ms and *σ* ∈ (10^−2^,10^0^) ms). Most of the performance data Δ distributed to form a plateau in the *T* and *σ* space (Figure 6D). In this plateau region, we defined *Q*=9 points corresponding to the combinations *C_σ,T_* = (*σ, T*), with *T* =2.25 ms, 3.5 ms, 4.5 ms, and *σ* =0.013 ms, 0.1 ms, 0.63 ms and we used them to reconstruct the connectivity matrix based on the threshold frequency *f_jk_* as formally described in the previous section.

The network model we adopted was used to generate spiking data useful to test and develop the connectivity algorithm based on analysis of correlations. However, this model was not intended to be representative of real biological neurons and does not reproduce all the specific features of electrical recordings from *in vitro* neuronal cultures. As a result, the range of parameters selected for the simulated case did not necessarily match the one for the real case and was later adapted to the data of recorded *in vitro* neuronal networks. However, the same parameters showed very high reproducibility in repeated experiments both with simulated networks and iPSC-derived neuronal systems. Scaling performances will be discussed later in the manuscript; however, *T*, *σ* and *ϵ* were not affected by the network size.

### Electrophysiological characterization of hiPSC-derived networks

Methods for generating cortical neurons from hiPSC, analyzing the composition of the resulting cell population, and culturing on MEA are described in the Supplementary Material. hiPSC-derived cortical neurons plated on 48-wells format MEA plates were recorded every week. Recordings were acquired with the Maestro recording system and Axion Integrated Studio (Axion Biosystems). A butterworth band-pass (10-2500Hz) filter and adaptive threshold spike detector set to 5.5X standard deviations were applied to the raw data. Raster plots of neuronal spiking activity were generated using Axion Neural Metrics Tool, and the data were analyzed using Excel (Microsoft) and GraphPad Prism version 7.00 (GraphPad Software) (Figures S3 and S4).

We used routines implemented in Matlab to analyze the electrophysiological recordings and identify the active neurons in the plate. In MEA recordings, the same electrode can record the activity of multiple neurons (Figure S5). However, identification of spikes corresponding to different neurons (Figure S6) is crucial to interpret electrophysiological recordings, especially in connectivity studies and analyses of causal dynamics in networks [52]. Therefore, we used spike sorting to group spikes with similar shape into different clusters, each corresponding to a different unit (neuron). This allowed us to isolate the activity of a few units per electrode, which resulted in the reconstruction of the activity of multiple detected neurons in the MEA well (Figure 8A). To this end, we adopted a commonly used spike sorting approach where principal component analysis (PCA) was used to extract similar features in the recorded spikes and clustering allowed us to group spikes with the same profile. We used a *k* - means clustering approach consisting of partitioning *n* observations into *k* clusters in which each observation belonged to the cluster with the nearest mean. The clustering algorithm started with a pre-defined *k* = 2. This value was then automatically updated to the best *k* estimate based on observations on the explained variance in PCA. The clustering was repeated 20 times using new initial cluster centroid positions (no change was observed for more than 20 replicates). The final output was the solution with lowest within-cluster sums of point-to-centroid distances. In a few cases, the clustering approach did not perform efficiently and generated either too many or too few clusters which were detected by observing the increased number of outliers in the spike sorting output. In these critical cases, a visual test was performed to identify the correct clusters.

### Connectivity analysis in hiPSC-derived cultures

We used our algorithm to estimate the connectivity in recorded iPSC-derived neuronal networks at week 1, 2, 3, and 4 (Figure 8). We selected a range for the parameter *T* based on considerations of the mean propagation delays between synaptically connected neurons. While physiologically many variables contribute to neuronal delays, a rough indication of the delay in synaptically connected cortical neurons was estimated to be 6-14 ms [20]. We estimated similar conduction velocities and latency values in a previous study performed on basket and pyramidal neurons from the rat neocortex [50]. Given that monolayer networks of human iPSC-derived neurons may have temporal properties that differ from those observed in intact brain networks, we decided to avoid neglecting correlations that could negatively bias the inference performance by considering a larger temporal window (−22, 22) ms. The reconstruction analysis was then performed with *T* ∈ 15, 22 ms to include the propagation delays measured in cortical neurons (6 - 14 ms) and over-estimated values to limit over-filtering. For *σ*, observations on the recordings and the level of smoothing required for high-frequency noise removal led us to select *σ* ∈ (0.3, 0.8) ms. We considered a fixed value of 3 ms equal to the average width of the correlation peaks. Within this range of parameters, we then selected *Q* = 9 points corresponding to the combinations *C_σ,T_* = (*σ, T*), with *T* =20 ms, 17.5 ms, 16 ms, and *σ* =0.4 ms, 0.55 ms, 0.7 ms.

We used a graph-based connectivity analysis to estimate developing connections between neurons. A common graph theory approach to measure the level of connectivity in a network is to investigate its centrality, which can be described as the capacity of a node to influence, or be influenced by, other system elements by virtue of its connection topology [43]. The simplest and most commonly used measure of centrality is node degree, which is the number of connections attached to a node. We calculated the average in-degree and out-degree of the vertexes of each directed graph corresponding to a reconstructed network by calculating all incoming and outgoing connections for each network vertex and then averaging over the total number of vertexes per network. The statistics was then extended to 20 different wells. A node scoring highly on a given centrality measure can be considered a hub. Here, we quantified the number of network hubs at each time point (from 1^*st*^ to 4^*th*^ week) by computing the centrality of the graph and ranking the vertexes based on the number of incoming connections. The most important vertexes were defined network hubs. We averaged this number over 20 reconstructed networks. Finally, we studied the integration and segregation properties of the networks [64] by computing the average Mean Path Length and Mean Cluster Coefficient (*CCo*). We calculated the Mean Path Length as the shortest path distance of all vertex pairs. Infinite (absent) connections between neurons were not considered in the calculation. With the use of Matlab routines, we also computed the *CCo* as the fraction of triangles around a node, which is equivalent to the fraction of nodes neighbors that are neighbors of each other [40].

## 3. Results

### Numerical results

To evaluate the performances of our method, we built a numerical model by designing and simulating an integrate-and-fire neuronal network mimicking the activity of *in vitro* cultures of neurons (see Supplementary Material). We studied *TP/N_c_*, *FP/N_c_* and Δ as function of the discrimination threshold *d* (Eq. 9) (Figure 7A). We note that, at a high discrimination threshold, while *TP/N_c_* remained roughly constant (the same true positives were always detected, for all points), *FP/N_c_* decreased towards very small percentages, smaller than the percentage corresponding to any of the *p* points. This is not surprising because the TP connections, which in most cases point to the interactions between the same pairs of neurons, were always observed for all points (*f_jk_* ≈ 1, *f_jk_* is the frequency for each connection in the connectivity matrix - see Eq. 9). On the other hand, the false positives are just fluctuations of the algorithm as new false positive connections pointing every time to a different pair of neurons were constantly generated by the algorithm for all points *p*. As a result, the same false positive connections observed for a given pair of neurons are very unlikely detected by many different points, and at high frequency they are filtered out.

**Figure 7:**
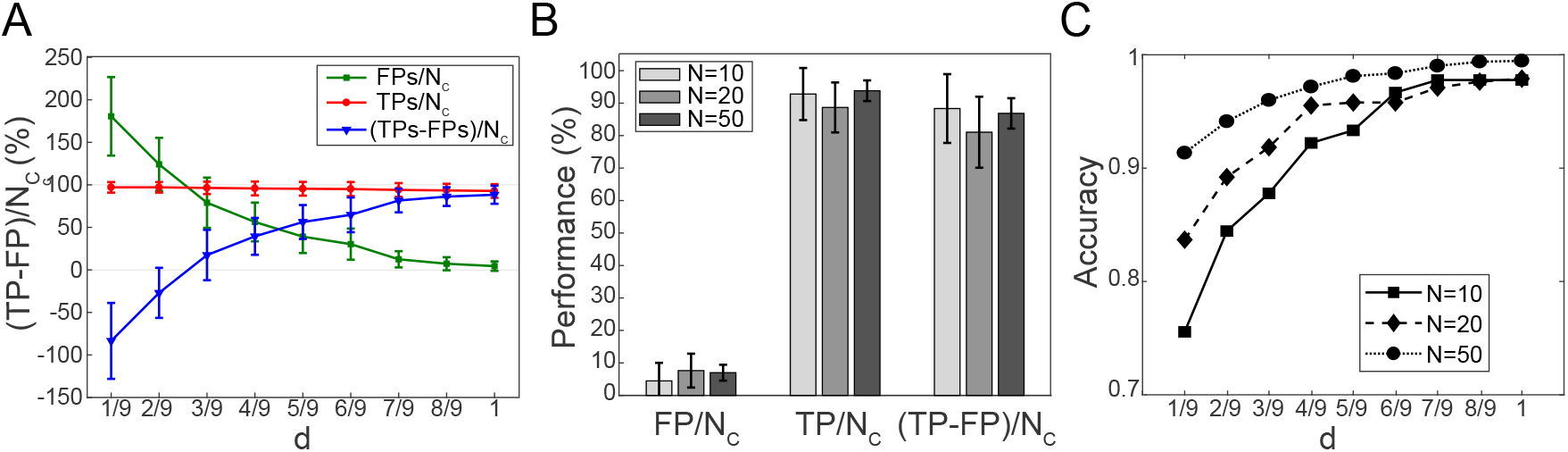
Method validation and performance evaluation. (A) Average (N=20) performances computed at different discrimination threshold *d*. The TPs remain roughly constant. On the other hand, the FPs decrease and fluctuate around very small percentage. The algorithm filters out the FPs that fluctuate at high frequency reaching best performances (85%) at *d* = 1. (B) Analysis of scaling properties. An average (N=20) number of *T P/N_c_*, *FP/N_c_*, and Δ was computed for 20 randomly generated networks with 10, 20, and 50 neurons, respectively. Very good performances are maintained constant for increasing network size. (C) Accuracy *ACC* indicator computed for different network sizes as a function of the discrimination threshold *d*. The error bars stand for the standard deviation of a dataset of 20 different networks.

For example, for two given points *p*_1_ and *p*_2_ the algorithm detected 15% and 20% FPs, respectively; however, most of the FPs detected by point *p*_1_ did not correspond to the ones detected by point *p*_2_. On the other hand, the same two points *p*_1_ and *p*_2_ detected the same TPs. Through filtering, the TPs detected by both points are preserved, all FPs are discarded, resulting in 2% of FPs remaining, a percentage that is smaller than any percentage associated with each of the points. Importantly, this result demonstrates empirically our mathematical framework and it highlights its robustness in this example application.

Δ data showed average performances of 88.3%, 88.7%, and 86.8% for varying network sizes of 10, 20 and 50 neurons, respectively, therefore demonstrating a very reliable detection of connections and exceptional scaling properties in this example (Figure 7B). Figure 7C shows the Accuracy of our method. It is interesting to observe how the accuracy data for *d* =1/9, i.e. prior to using the reconstruction algorithm based on the discriminator threshold *d* (Figure 7C, *d*=1/9), show performances already higher than 75% for a network of 10 neurons, and even higher for larger sizes, thus exceeding the ones obtained before standard thresholding in published connectivity methods [47]. The complete reconstruction algorithm led to a computational accuracy close to 100% for all network sizes due to the further statistical pruning of false positive connections (Figure 7C, *d* =1).

It is worth noting that, compared to more conventional indicators, Δ is independent of the network’s connectivity. Since we made the hypothesis that each node in the network model has the same average connectivity (preserved in- and out-degree) which does not increase with the network’s size, we do not expect and we are not interested in seeing connectivity dependent variations. On the contrary, other metrics, such as the Receiving Operating Curve (ROC) or the Accuracy [49], largely change with the connectivity properties of the network. For comparisons, we calculated the Accuracy of the investigated network model. However, because the definition of this measure is based on the hypothesis that larger networks feature higher connectivity, our data overestimate the performances, especially for high network sizes (see the Discussion for further details on this).

### Measures of developing iPSC neural networks in vitro

We tested our connectivity algorithm on networks of human iPSC-derived neurons cultured on 48-well MEA plates (*Axion Biosystem*). The activity was recorded periodically over four weeks (Figure S3) to investigate network development. In parallel, immunofluorescent analyses (Figure S4) of the composition of the neuronal population showed a mix of excitatory and inhibitory neurons, as well as astrocytes, resembling the physiological composition of an *in-vivo* neural network.

The data show an increase in the level of activity as a function of the developmental phase, as well as the appearance of repetitive firing patterns and the formation of well-organized network bursts (Figure S3).

We used our algorithm to estimate the connectivity in the recorded iPSC-derived neural networks at week 1, 2, 3, and 4 (Figure 8B). Based on the position of each electrode, we estimated the localization of the recorded neurons. The neurons have been randomly distributed around their corresponding recording electrode, within a radius of 50 *μ*m, as this is the expected sensitivity range of MEA electrodes. Figure 8C reports the corresponding graph-based schema of the reconstructed adjacency matrices in B, for week 1 and 4, respectively. Both descriptions clearly demonstrate maturation of the network and increase in overall connectivity, as well as the formation of a few highly connected sub-networks within the culture.

**Figure 8:**
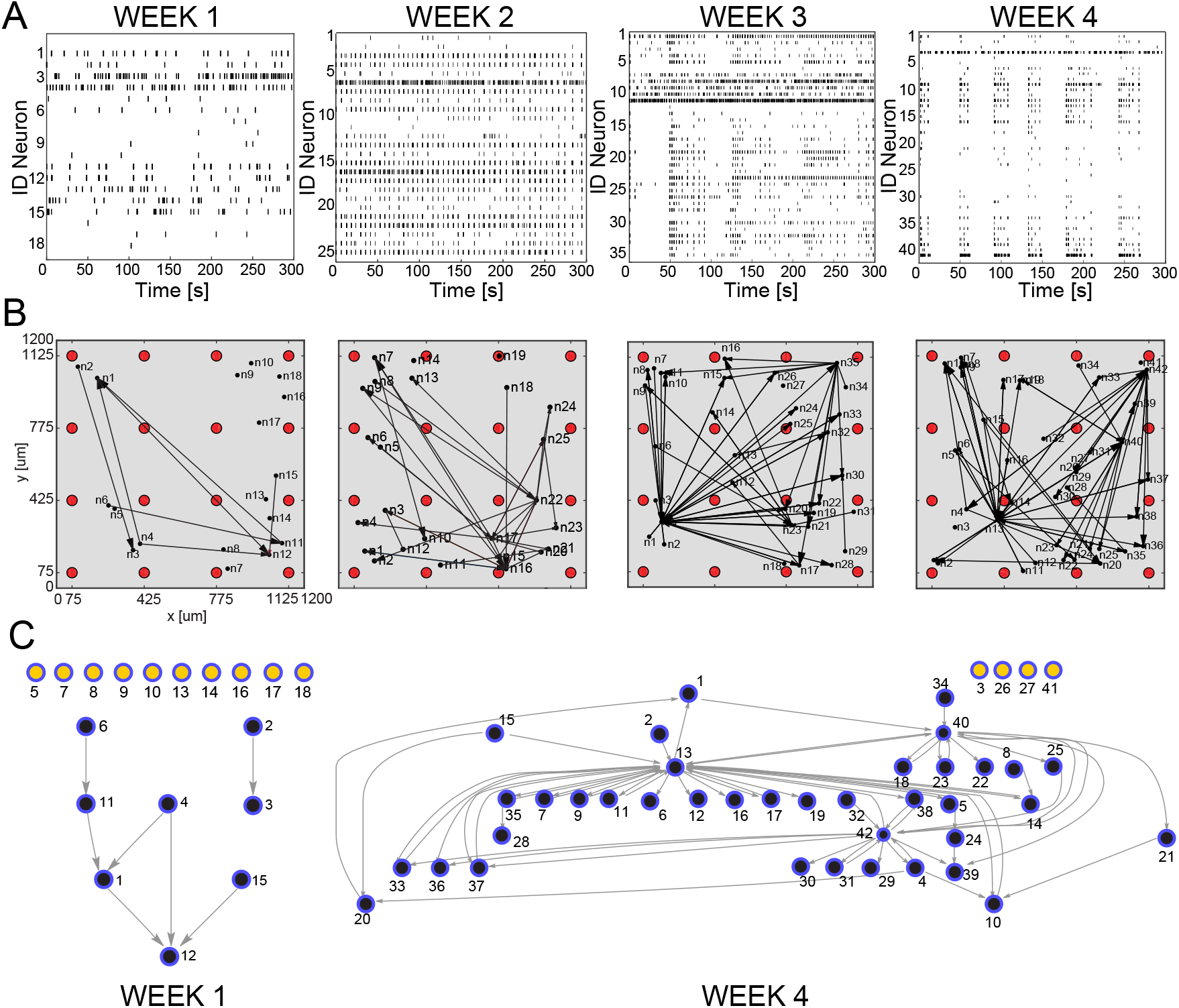
Connectivity reconstruction from neural recordings of developing hiPSC-derived neurons. (A) Raster plots of the same MEA well at different time points during development: week 1, 2, 3, and 4 after plating. The four panels shows spiking signals from individual neurons (rows) obtained through spike sorting, PCA and *k*-means clustering of 300 sec recordings. The culture develops complex network features: from weakly active and randomly organized (individual spiking events), to very active and fully organized (network bursts). (B) Estimated effective connectivity of the developing culture whose activity is described in panel A. Each visual map consists of a 1.2 x 1.2 mm MEA plate (gray area), a 4-by-4 array of micro-electrodes (red circles) and the estimated directed connections (black arrows). The active neurons are represented by black dots; they are randomly distributed around their corresponding sensing electrode within a radius of 50 *μ*m. (C) Directed graph relative to the culture at week 1 and week 4. The connectivity is equivalent to the one visualized in B but, for clarity and consistency with the main text, links between neurons are directed edges (arrows), active neurons are network nodes (blue: connected; yellow: independent). Given a MEA well and a specific time point, indexes refer to active neurons with recorded activity reported in (A). Note, neurons mapped at week 1 do not correspond to neurons mapped in the following weeks, although they are indicated with the same indexes.

To generate a more quantitative estimation of these network features, we used a graph-theory based analysis (Figure 9). The larger number of active neurons and detected links in most developed networks is in accordance with the increasing levels of activity of the culture at later time points (Figure 9A). We observed that during spontaneous activity, the in- and out-degrees for different neurons were almost equally distributed among the active neurons, and showed an increasing trend as a function of the culture’s age. Moreover, we observed formation of a specific network topology characterized by a connectivity highly centralized around a few super-hubs reciprocally linked to neighboring cells (large average degree), and a few remaining, poorly connected neurons (small average degree) (Figure 9B). These data confirm what is visually described in the connectivity maps and correlate with the network burst activity in the way the connectivity topology changes from random and unorganized, with most of the neurons isolated and poorly connected, to extremely structured and centered around a few hubs, a topology that becomes more evident with maturation. We also studied the integration and segregation properties of cultured human iPSC-derived neural networks (9C). The Mean Path Length is fairly constant and does not change for increasing numbers of new connections. On the other hand, the Mean Cluster Coefficient (CCo) is low and decreases with the level of maturation of the network. This feature is indicative of more favored segregation versus integration, with formation of highly integrated network units (hubs), similarly to observations reported by Livesey and colleagues [30] using rabies-virus-based trans-synaptic tracings of hiPSC-derived neuronal networks.

**Figure 9:**
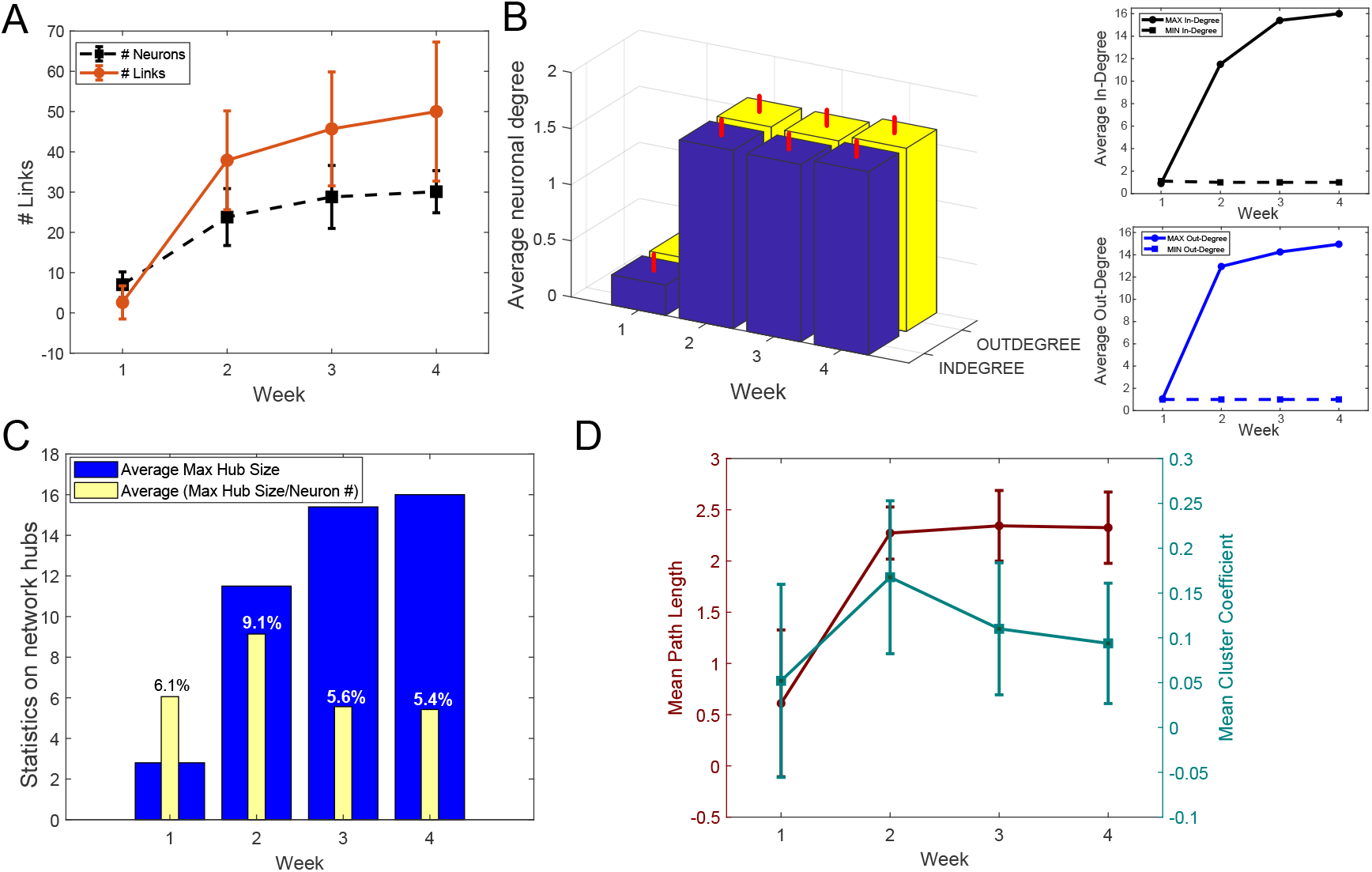
Graph-theory based analysis of network connectivity in developing hiPSC-derived neuronal-network. (A) Average number (N=20) of active neurons (black) and detected connections (red) in MEA wells recorded at week 1, 2, 3, and 4 after plating. The error bars represent the standard deviations for a dataset of 20 different MEA wells. (B) (Left panel) Statistics (N=20) on the neuronal in-(number of input connections) (blue) and out-degree (number of output connections) (yellow). The vertical red bars stand for the standard deviations of in- and out-degree data. (Right panel) Average maximum (continuous line) and minimum (dashed line) in-(top) and out-(bottom) degree. The number of in- and out-degree is equally distributed with equivalent raising behavior as a function of the cultures’ developmental stage. (C) Statistics on network hubs as a function of the culture’s age. In blue, average (N=20) maximum hub size. We used the in-degree centrality measures for hub detection and characterization. In yellow, percentage of neurons that function as network hubs relative to the total number of active neurons in the well (value averaged over 20 wells). More mature networks include few super-hubs (∼5% of the total number of neurons) with increase in size as demonstrated by the raising number of incoming connections at week 3 and 4. (D) Characterization of network segregation and integration properties. Values in the graph correspond to the mean Path Length (PL) and mean Clustering Coefficient (CCo) calculated for each well at week 1, 2,3, and 4 after plating and then averaged over 20 wells. The error bars correspond to the standard deviation of 20 different measures. The mean PL corresponds to the average shortest path length in the networks. Infinite (absent) connections between neurons were not considered in its calculation. The PL is very low for highly immature cultures where only sparse activity from individual neurons was mainly registered. It increases as soon as connections are formed (week 2) but no longer changes with the number of new connections (week 2, 3, 4). The CCo is low and decreases with the maturation of the network as indication of favored segregation versus integration with increasing number of independent but highly-integrated network units (hubs).

## Discussion

In this work, we demonstrated a new model-free based approach to infer effective - active, direct and causal - connections from *in vitro* neuronal networks recorded on MEAs. Our algorithm offers several fundamental differences resulting in critical advancements compared to the state-of-the-art connectivity techniques, including correlation and transfer entropy variants [14, 47]. To the best of our knowledge, model-free connectivity inference techniques are not able to reconstruct the effective connectivity of a recorded neuronal network because they are either missing the causality of signaling or include confounding apparent connections (common input or common output) and multi-neuron pathways. Here, we defined a fundamental mathematical rule that decomposes the misleading temporal information contained in the network’s temporal correlations into a set of direct, causal dependencies between the circuit’s neurons via selective identification and elimination of false positive connections (Figure 2).

Our method does not require any post-inference thresholding and it only depends on three fundamental parameters - *T*, *σ* and *ϵ* - which can be easily obtained as explained in the text, and whose choice will be automated in future software release.

Evaluation of the performances on synthetic data from simulated random networks of neurons demonstrated a number of critical improvements compared to published algorithms by showing inference of causal, uni- and bi-directional connectivity directly from MEA data with a computational accuracy close to 1 and an excellent rate of rejection of false positive connections (Figure 6).

In terms of scalability, our algorithm matches the scalability properties of most of the state-of-the-art connectivity methods which, being based on pairwise statistical and correlation studies, scale with a practical computational complexity of the order of *O*(*n*^2^), where *n* is the number of neurons. More specifically, we have three main routines whose complexity should be assessed: the computation of pairwise correlations, the peak detection algorithm, and the detection of false positive connections. In order to do this, let *S* be the number of time samples of a recording for each neuron. The bare complexity of the correlation computation is *O*(*n*^2^*Slog*(*S*)) since it involves Fast Fourier Transform evaluations. The complexity of the peak detection algorithm, on the other hand, is *O*(*n*^2^*S*). Finally, the computational complexity of the brute force detection of false positive connections is *O*(*n*^3^). However, a closer look at these estimates shows that the computation of the correlation is the practical leading term. In fact, it has a larger pre-factor with respect to the peak detection which results from the several algorithm steps included in the Fast Fourier Transforms. Moreover, the peak detection can be restricted to just a much smaller time interval thus reducing the *O*(*n*^2^*S*) complexity to *O*(*n*^2^*S*′) with *S*′ « *S*. Finally, the computation of correlation triangles is negligible with respect to the computation of the correlations for practical cases. In fact, if we consider a standard neuronal recording, this typically has more than 10^6^ time samples. Since the computational complexity of all correlations is *O*(*n*^2^*Slog*(*S*)), unless we are dealing with more than millions of neurons, this is usually much larger than *n*^3^ three terms floating point operations needed to assess the false positive connections. Moreover, with a more sophisticated approach based on dynamic programming we could compute triangles with a reduction of the *O*(*n*^3^) complexity to *O*(*n*^2^) for practical cases. However, the dynamic programming implementation details is out of the scope of this paper, even if a basic implementation is included in our software (https://github.com/fpuppo/ECRtools.git). Comparison with other techniques such as partial correlation, matrix deconvolution or deep learning, which we found to be the only approaches able to distinguish between apparent connectivity (see Figure 1), shows usually higher computational complexity. For example, partial correlation yields a complexity of *O*(*n*^3^*S*).

Importantly, in this work we did not attempt to simulate and analyze large artificial neural networks where the number of nodes and connections aim to mimic the massive synaptic connections present in the brain. We simulated sparse networks instead, having connectivity *N_c_* < *N_tot_*, and hypothesized average constant connectivity for larger network sizes. This simplified model was quite accurate in describing *in vitro* networks of biological neurons like the cultures of iPSC-derived neurons we investigated (see Figure S.4). Larger networks can be simulated and tested at higher computational expenses. Based on observations on the algorithm performances, we expect approximately constant performances for increasing number of neural nodes; on the other hand, we can only speculate on the inference properties in highly connected networks (*N_c_* ∼ *N_tot_*) because of increased difficulties in properly modeling their simulated activity, as well as the anticipated increases in computational power required for processing these analyses. We expect decaying performance for networks of very high complexity due to many overlapping correlation effects from multiple cells, a problem that will require further investigation and will be addressed in future studies.

As a direct example, we have used the spontaneous firing of cultured networks of iPSC-derived neurons to reconstruct the corresponding connectivity maps as a function of the network developmental stage (Figure 9). The estimated connectivity (Figure 8) combined with quantitative analyses, such as graph-theory approaches (Figure 9), enabled us to describe the developmental progress of the cultures, thus demonstrating the capability of detecting basic neuronal features such as the increased synaptic connectivity and the formation of few, highly connected network hubs. These latter are both indexes of more mature neuronal circuits and agree well with higher spiking frequencies and network burst generation observed in mature neuronal cultures [71].

To estimate causal and direct connections in iPSC-derived networks, we relied on conventional spike sorting techniques in order to decompose MEA recordings into spiking activities corresponding to individual neurons. However, it is worth mentioning that despite its long history and substantial literature, spike sorting remains one of the most important and most challenging data analysis problems in neurophysiology. Spike sorting based on PCA and *k* - means clustering is one of the most accepted methods in part due to its ease of implementation. However, a number of different approaches have been proposed over the years [9]. Although we recognize that the accuracy of the spike sorting procedure critically affects all subsequent analysis [52], further treatment of this issue goes beyond the scope of our paper. Indeed, testing of performance and validation of the connectivity algorithm we introduce in this work requires knowledge of the ground truth (i.e. knowing the identity of the neurons generating each detected spikes) which is only really possible in synthetic models. When applied to biological networks, our reconstruction method assumes that the spike sorting is accurate enough to separate the most prominent contributions in the activity of the network. However, the higher the accuracy of the adopted spike sorting procedure, the more precise the estimation of causal activity in the network will be. In other words, our network reconstruction methods scale with continued improvements in spike sorting.

Furthermore, since our algorithm fully relies on neural recordings to estimate the effective connectivity of the network, the connectivity inference also depends on the recording technology. The quality of any connectivity method, including the one we introduce here, can only be as good as the available measured empirical data. MEAs devices typically available in laboratories have a low density of electrodes (typically 8×8 or 64×64 array size) which result in low spatial resolution of the sensitive area (spacing between the electrodes) and low spatial granularity in the recordings. The main consequence is reduced detection of active neurons, which affects the accuracy of reconstructing the real networks activity. This is valid for all functional connectivity methods. In our approach, unrecorded units and associated undetected correlation patterns may produce a reduction in the number of computed correlation triangles, which in turn may result in reduced filtering capabilities and unidentified false positive connections. As other technologies from the Brain Initiative [31] and related efforts, such as the Human Brain Project in the EU, become available online and are validated, the methods we develop here will be in a position to take immediate advantage of them. For example, high density MEAs (HDMEAs) that include tens of thousands of electrodes at cellular and subcellular resolution [73, 80, 38, 7], and complementary voltage and calcium imaging approaches [26, 41, 3, 27] that provide useful information about the localization and classification of recorded neurons, offer optimized acquisition settings and will greatly improve the resolution and accuracy of our approach. To this end, our group is actively engaged in exploring novel recording technologies that can further validate and improve the connectivity algorithm we discuss here. Complementary to high-resolution CMOS-MEAs, high density arrays of individually addressable nanowires represent a viable approach to increase spatial resolution, scalability and sensitivity of the recordings. For example, the system by [32] can detect sub-threshold events, thus enabling the recording of miniature post-synaptic currents that can potentially be involved in structuring network connectivity.

Importantly, our method is scalable and can be generalized to any kind of network, thus allowing the user to target different problems in intact neurons, including synthetic models as well as *in vitro* and *in vivo* systems. Of particular relevance and as an example of an on-going effort to address these issues, our group is exploring the use of technologies to analyze electrophysiological recordings from 3D cultures. We are assessing different techniques for high-density 3D electrophysiological characterization of human derived cortical organoids [71]. The application of the methods we discuss here to the connectivity analysis of 3D neural structures will allow us to test a variety of hypotheses about the development of 3D neuronal networks, dynamics and changes that may occur under external perturbation or in disease.

A limitation of the system is the inability to detect sub-threshold events due to the extracellular nature of the multi electrode array recording, and thus to investigate the potential relationship between miniature excitatory and inhibitory post-synaptic currents and connectivity. Recently, the application of nanotechnology on neuronal electrophysiology has brought about a promising solution to overcome this limitation and to fabricate devices that are capable of detecting sub-threshold potential [32, 77, 81]. Although substantial engineering issues remain before the potential of nano neural interfaces can be fully exploited [79], the future application of our algorithm to more sensitive recordings appear to be a promising approach for a more accurate understanding of network connectivity as a consequence of synaptic and extra-synaptic inputs as well as subthreshold potentials in both physiological and pathological conditions.

Finally, in this work, we have estimated the connectivity matrix of a sub-network of excitatory links which have been described as the strongest recurrent links in the neuronal culture, major determinants of spontaneous activity [67]. However, it is increasingly clear that inhibitory connections play essential roles in neural dynamics. Therefore further work must also be done to extend this approach to the explicit identification of inhibitory inputs and their role. Future improvement will include integration of cross-correlation approaches previously investigated by other groups for the detection of inhibitory effects [46], and extension of the correlation triangle technique to account for inhibition. A similar problem exists to detect excitation when there is strong tonic excitation close to saturation or persistent bursting. In the bursting regime, a functional reconstruction typically results in highly clustered connectivity due to the highly synchronized firing of large communities of neurons that appear to be all connected even though no direct synaptic connectivity exists [67]. In our method, the bursting regime can induce an increase in the number of computed correlation triangles with potential underestimation of false positive connections due to the large number of apparent correlations between synchronized cells. However, since the Izhikevich model we used for validation includes also bursting neurons, we think that the method here proposed can already behave quite well under bursting conditions too. Nevertheless, we are currently investigating a novel approach to improve the accuracy of estimation of connectivity between bursting neurons; however this problem remains a matter for future work.

## 4. Conclusions

We have presented an innovative approach to map the effective connectivity of neural networks from multielectrode array data. The tool that we have developed offers critical improvements over available methods for estimating functional connectivity. Notably, our connectivity algorithm succeeds in detecting direct connections between neurons through a mathematically rigorous selection scheme that distinguishes between apparent or non-direct links and direct ones, therefore enabling inference of directed causal relationships between connected neurons. In addition, it has good scaling capabilities and can be further generalized to any kind of network, thus allowing to target different problems in intact neurons, synthetic models as well as *in vitro* and *in vivo* systems. As novel electrophysiology technologies come online and are validated, the methods we presented here will be in an immediate position to take advantage of them, resulting in fundamental improvements in spatial resolution and reconstruction accuracy. Furthermore, our algorithm can be further extended, improved, and possibly integrated with already in-use techniques to overcome important limitations such as the detection of inhibitory connections and the inference of effective connectivity in the bursting regime.

Importantly, spatio-temporal information is implicitly contained in the estimated connectivity and delay map; we expect therefore that, when used in combination with novel computational methodologies [60, 50, 6], our method will help reveal more fundamental network properties crucial to the understanding of the relationships between network topology, dynamic signaling and network functions in healthy and disease models. Furthermore, it will be broadly applicable to experimental techniques for neural activation and recording, increasing its utility for the analyses of spontaneous neural activity patterns, as well as neuronal responses to pharmacological perturbations and electrical and optogenetic stimulations [70, 26, 38, 41, 3].

## Acknowledgments

The authors thank Fabio L. Traversa for fruitful and constructive discussions on the theoretical framework. This work was funded by a Swiss National Science Foundation (SNSF) Mobility Fellowship (P2ELP2 168553) to FP, a NIMH U19MH106434 to AB, and in part by unrestricted funds to the Center for Engineered Natural Intelligence (CENI).

## Author contributions

F.P. conceived the original hypothesis and theoretical approach, with additional contributions by G.S‥ F.P., D.P., A.B. and G.S. conceived the project. F.P. developed and implemented the theory, organized the study and carried out all the computational analyses. D.P. and A.B. designed and organized the experimental studies. D.P. carried out all experiments and related analyses. All authors have contributed in interpreting the results and writing the manuscript.

## Materials and correspondence

Correspondence and request for materials should be addressed to G.S. and A.B.

## Supplementary materials

### Neuronal network model

We validated the connectivity method on spiking data generated via simulations of neural networks based on the Izhikevich’s model [1]. The original program was copied directly from his paper and modified to guarantee high levels of activity in the network as well as bursting like behavior similar to that registered in our experiments.

The voltage of each simulated cortical neuron are described by the coupled differential equations:

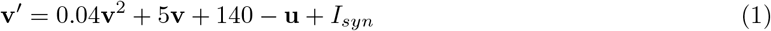

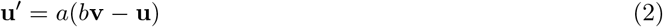

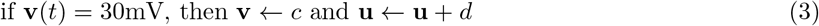

where **v**is the neuron’s voltage, **v**′ is the time derivative of the voltage, **u** is a recovery variable, **u’**is the time derivative of the recovery variable, *I_syn_* is the total synaptic input received by the neuron, and *a*, *b*, *c* and *d* are adjustable parameters that govern the firing behavior of the neuron. Here, the notation ← indicates that the variable **v** will be assigned the value of *c* if the conditional statement is true. The units of time are milliseconds and the units of voltage are mV. To simulate a network of regular spiking and bursting cells, we used (*a*, *b*, *c*, *d*) = (0.02, 0.2, −65, 8).

We used the variable *I_syn_* as excitatory input delivered at random times to all neurons, or a selected set of them, and simulated as a Poisson process with a mean firing rate defined *ad-hoc* for each experiment. As approximation, all neurons were considered excitatory with a homogeneous synaptic weight of 0.5 mV. Figure S.1 shows an example of random network generated via the implemented model.

### Generation of in vitro neural networks of iPSC-derived neurons

All human stem cell culture was performed under approval from the Stem Cell Research Oversight (SCRO) panel at Sanford Burnham Prebys Medical Discovery Institute. Undifferentiated hiPSC [2] (provided by Drs. H. Song and G. Ming, University of Pennsylvania), were cultured on irradiated mouse embryonic fibroblasts in hPSC medium (medium composition described in ref. [3]). To expand, hiPSC were passaged weekly as small clusters of cells using the ethylenediaminetetraacetic acid (EDTA) solution Versene (Gibco—Thermo-Fisher Scientific) as described in [4]. Cortical neural progenitor cell (NPC) were differentiated from hiPSC as embryoid bodies as described in [2]. To differentiate to cortical neurons, NPCs were dissociated with Accutase (StemCell Technologies), plated at a density of 1 x 10^6^ cells/cm^2^ on a 60 mm tissue culture dish coated with 20 *μ*g/ml laminin from Engelbreth-Holm-Swarm murine sarcoma basement membrane (Sigma-Aldrich) and then cultured for 2 weeks in neural maintenance medium (medium composition described in [3]). After two weeks, neural maintenance medium was switched to a medium comprised of Neurobasal A (Gibco—Thermo-Fisher Scientific), 10% Knock-out serum replacement (Gibco—Thermo-Fisher Scientific) and 1% Penicillin streptomycin (Cellgro), and the culture was further differentiated for two more weeks. To prepare the differentiated culture for plating on imaging and MEA plates, cells were dissociated with Accutase (StemCell Technologies) for 30 minutes, strained through a 70 *μ*m cell strainer and then re-plated at a concentration of 150,000 cells/cm^2^ in Neurobasal A, 10% Knock-out serum replacement, 1% Penicillin streptomycin and 1 *μ*g/ml of laminin on a 35 mm tissue culture dish pre-coated with 20 *μ*g/ml laminin. After 1 day the cells were fed with the same medium plus 5 *μ*M cytosine arabinoside (Ara C) (Sigma-Aldrich). After 2 days of Ara C exposure, the cells were plated on 48 well MEA plates (16 electrodes/well) (Axion Biosystems) previously prepared following the manufacturers instructions, at a concentration of 75000 cells/well in a 10 *μ*l drop. In parallel, AraC treated hiPSC derived cortical neurons were plated in 384 well imaging plates (Poly-D-lysine treated, Biocoat, Corning) coated with laminin (20 *μ*l/well) for 1h at 37°C at a concentration of 5000 cells/well. Cells were maintained in a humidified 37°C incubator with 5% CO_2_, and 66% of the medium was exchanged every other day. One week after plating the medium was changed to Brainphys medium (Stemcell Technologies).

### Immunocytochemistry

Neurons on 384 well imaging plates were fixed in 4% paraformaldehyde (PFA) (Alfa Aesar Chemicals) for 10 minutes at room temperature at 2 and 4 weeks after plating. The cells were then washed 3 times in phosphate buffered saline (PBS) (Gibco—Thermo-Fisher Scientific), and then blocked in 5% donkey serum (Jackson ImmunoResearch) and permeabilzed in 0.1% Triton-X (Sigma-Aldrich) in PBS. Primary antibodies at the dilutions noted (Table 1) were incubated on the cells overnight at 4°C. The following day cells were washed 3 times in PBS, and the secondary antibodies (AlexaFluor, Molecular Probes) were added 1:500 in blocking buffer for 2 hours at room temperature. The secondary antibody was then washed several times in PBS and DAPI (Thermofisher, 1:2000 in PBS) was added for 30 minutes at room temperature. Images were acquired with the Opera Phenix High Content Screening System confocal microscope (Perkin Elmer).

### Tables and Supplementary figures

**Table 1:**
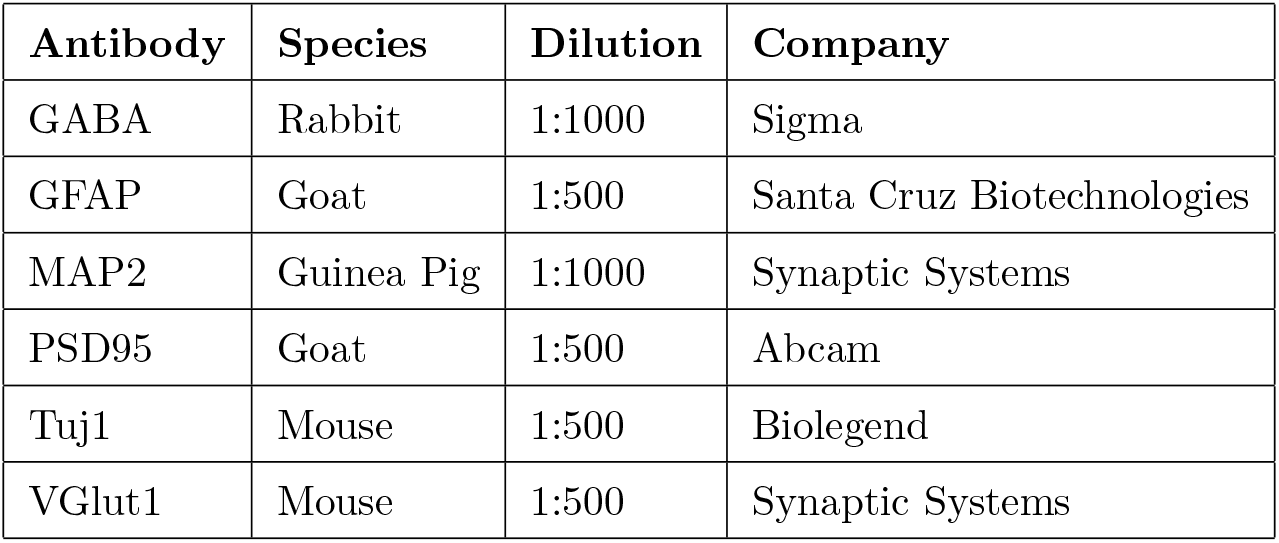
Antibody dilutions.

**Figure S.1:**
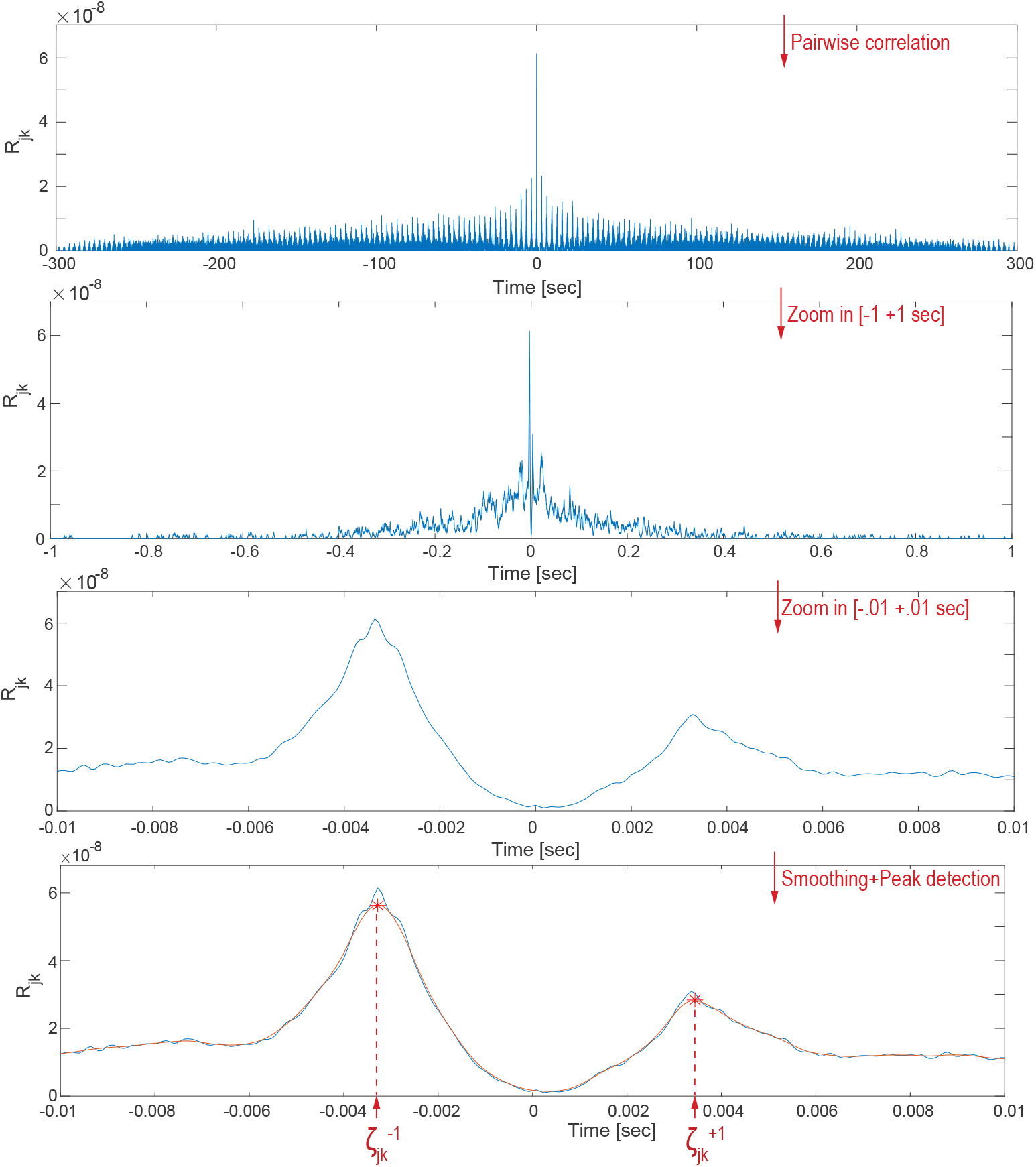
Correlation analysis and temporal delay detection. Correlation function computed on a pair of neurons (*j, k*) between their corresponding spiking signals *s_j_* and *s_k_*. Subsequent zooming is applied to show examples of correlation peaks. Detection of two correlation peaks is visually demonstrated in a arbitrarily defined temporal window *T* = (−10, 10) ms. Smoothing is applied to the signal through Gaussian filtering (orange line), and then followed by peak detection on the smoothed curve. The location of each correlation peak (red stars) with respect to the origin is a first approximation prediction of the temporal delay 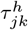 between the two neurons firing. Peaks detected on the positive or negative side indicate whether neuron *j* fires before or after neuron *k* and is therefore indicative of the directionality of communication flow between *j* nd *k*. The amplitude of the peak is calculated on the *R_jk_* function in correspondence of the peak location.

**Figure S.2:**
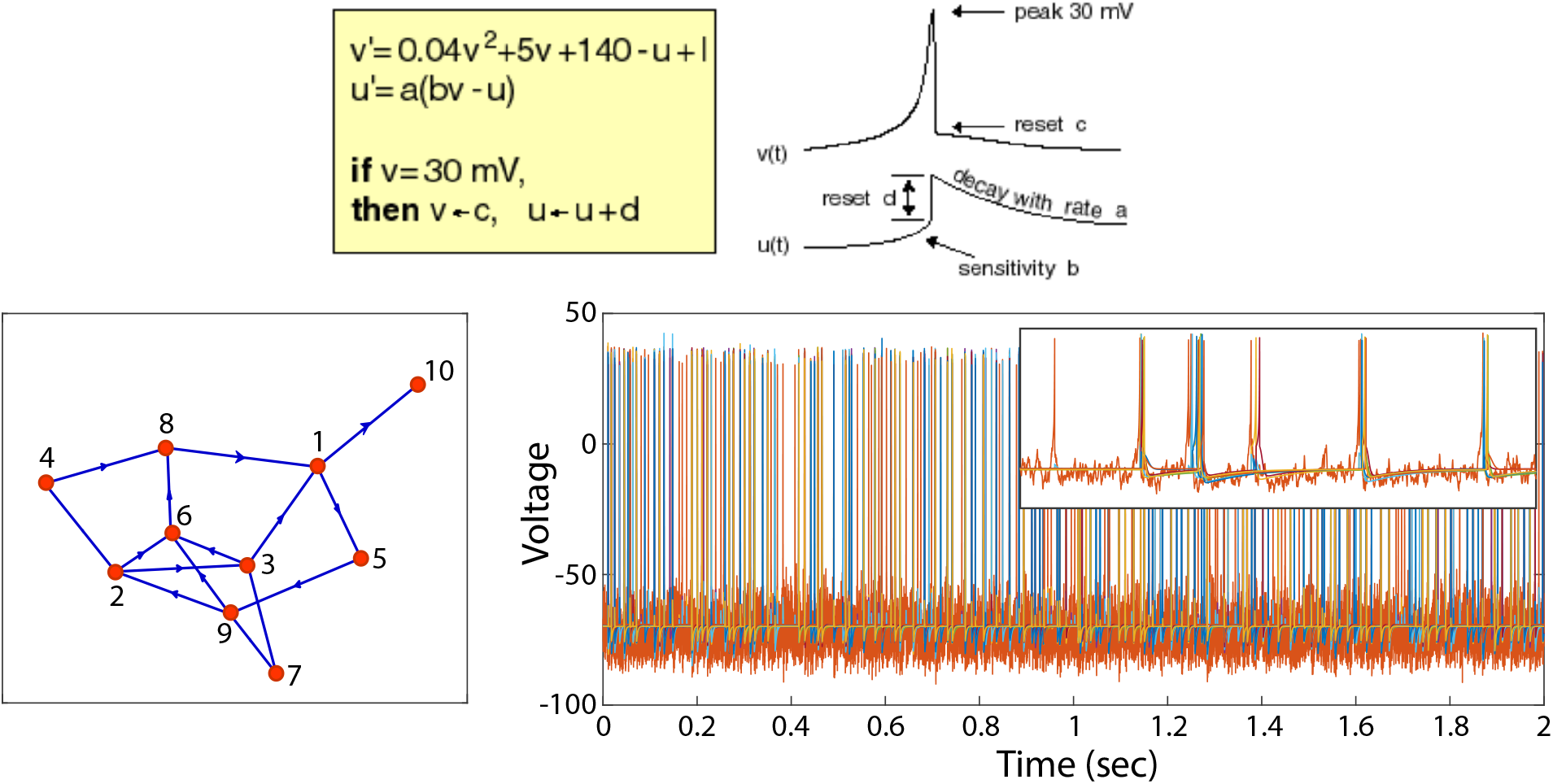
Neuronal network model. (A) Izhikevich model of spiking neurons. On the left, the equations of the model. On the right, visual description of the model’s parameters. (reprinted from [1]). (B) Directed graph (blue: edges; red: network’s nodes) corresponding to an example random network of 10 neurons. (C) 20 seconds long simulation of the 10-neurons network displayed in B. Different colors stand for different active neurons. Inset: 2 sec zoomed view on the spiking activity.

**Figure S.3:**
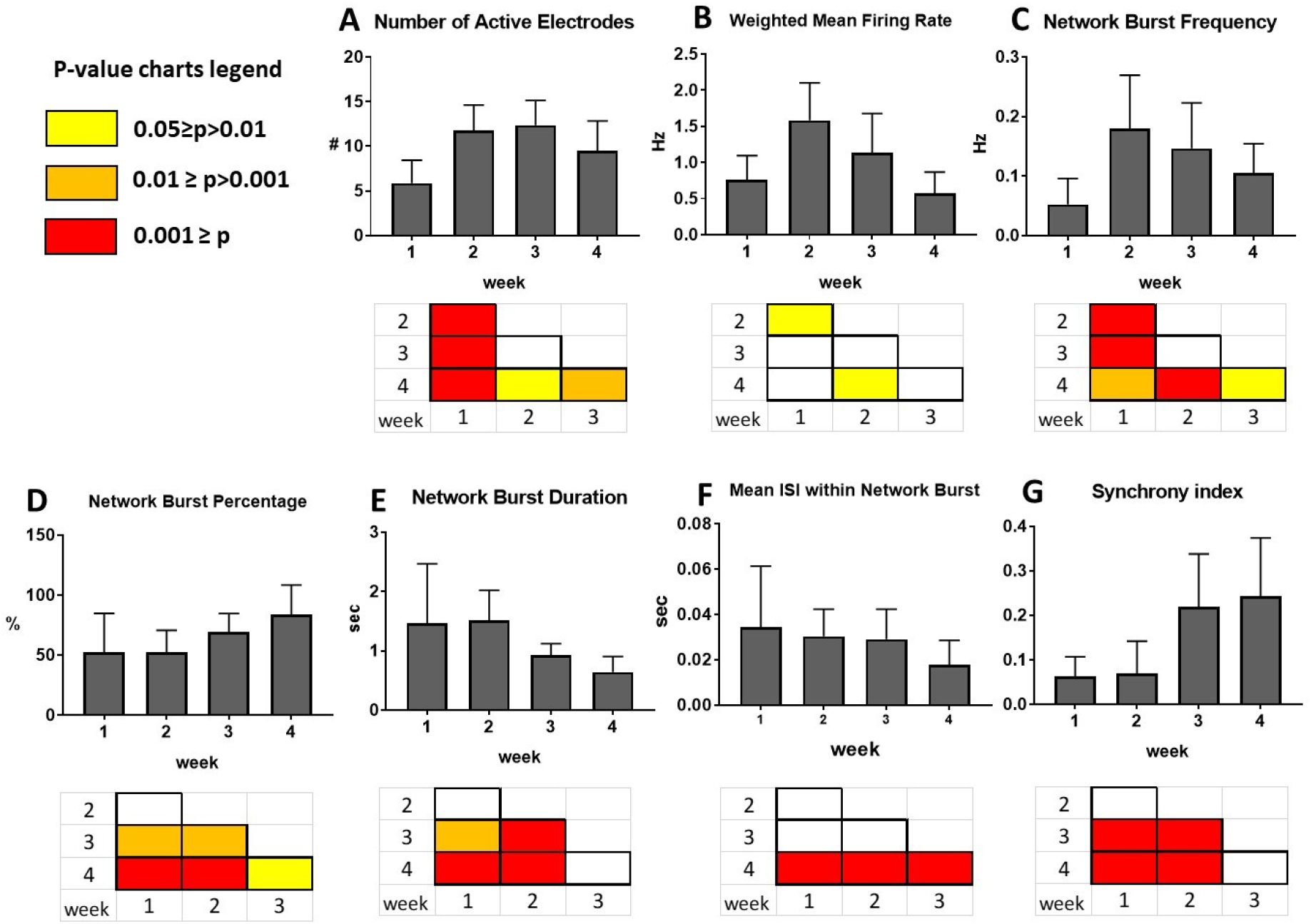
Development of neuronal activity and network characteristics over 4 weeks. Values are represented as average over n=24 wells and error bars indicate standard deviations. Statistical significance as heatmaps of p-values from paired t-test is shown for pairwise combinations of each week below each graph. See Methods for definition of parameters. (A) Number of Active Electrodes per well over 16 electrodes, showing a good number of electrodes recording activity over time, as well as an increase between the first and the second week. (B) The Weighted Mean Firing Rate shows an increase in neuronal activity between week 1 and week 2, while the activity decreases by week 4, as the network organization increases. (C) The Network Bursts Frequency sharply increases between week 1 and 2, when the network is formed, and then slightly decreases by week 4. (D) The Network Bursts Percentage increases over time, indicating the realization of a stable connected neural network and a reduction in sporadic extra-network firing. (E) The Network Burst Duration decreases at week 3 and 4 because of an increase in the spike frequency within a Network Burst, as shown from the decrease of Mean Inter Spike Interval (ISI) within Network Burst in panel (F), while the number of spikes per burst and the number of spikes per network burst remain constant (data not shown). (G) The Synchrony Index increases from week 2 to week 3, indicating increased participation of electrodes in synchronous network bursting.

**Figure S.4:**
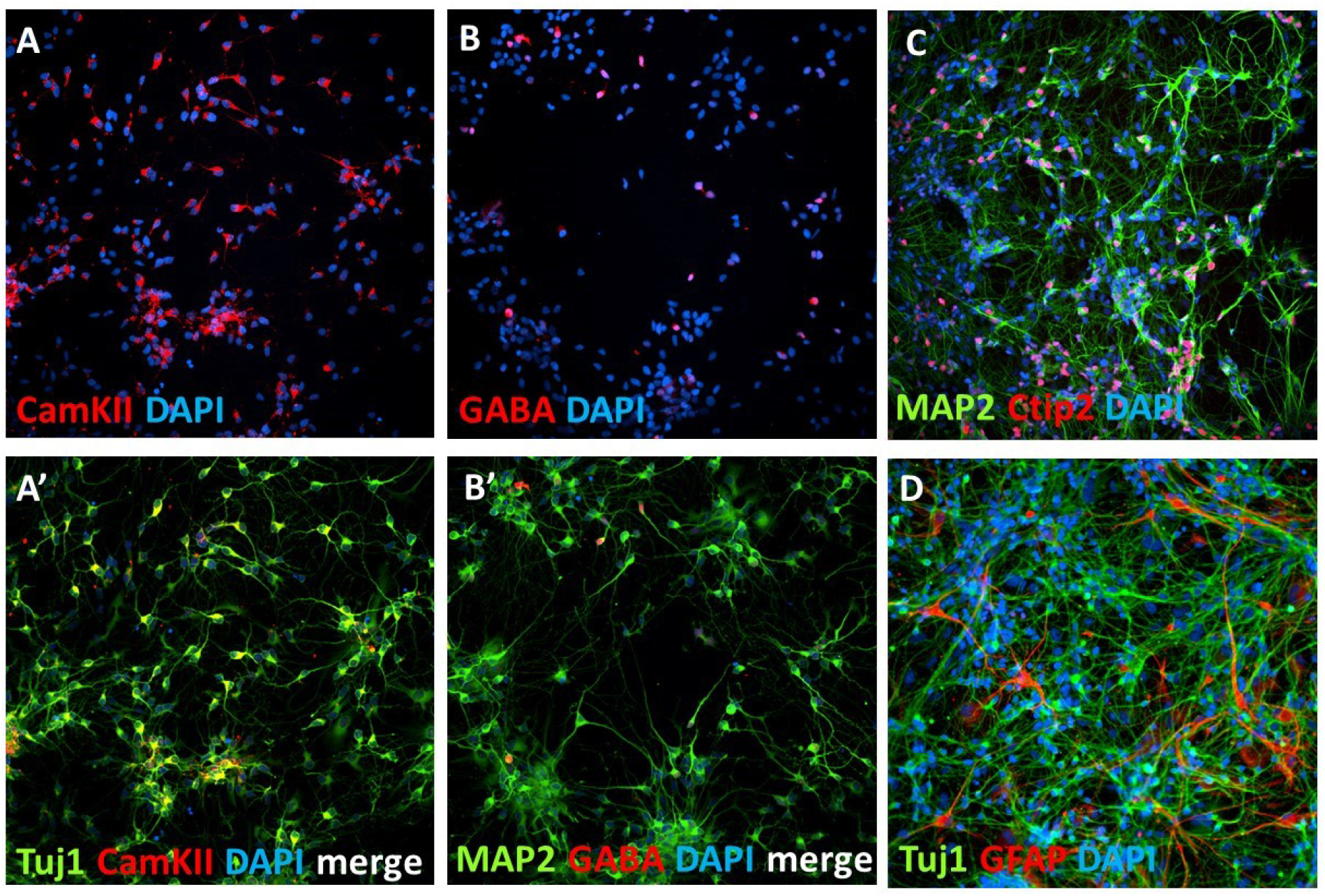
Characterization of human iPSC-derived cortical like neuronal culture. Neuronal culture differentiated for 4 weeks from NPC stage, re-plated on 384 well imaging plates, and then differentiated for an additional 1 week (panels A and B), 3 weeks (C) or 4 weeks (D). The percentage of positive cells are expressed as Average ± standard deviation. (A) and (A’) show immunostainings for CamKII (red), a glutamatergic marker and Tuj1 (green), a neuronal marker. The percentage of TUJ1 positive neurons is 66.7%±6.8. The percentage of TUJ1 positive neurons that are co-positive for CamKII is 89.2%±6.5 (with an average of 2827±342 cells/well, n=4 wells, 9 fields per well). (B) and (B’) show immunostainings for GABA (red), an inhibitory neuron marker and MAP2 (green), a dendritic marker. The percentage of TUJ1 and GABA co-positive GABAergic neurons is 8.12%±1.9 (with an average of 2377±335 cells/well, n=4 wells, 9 fields per well). (C) Immunostaining showing cells positive to CTIP2 (red), a transcriptional factor expressed by deep layer (V and VI) neurons (percentage of positive CTIP2 and MAP2 co-positive neurons is 41.7%±4.3, with an average of 3895±875 cells/well, n=4 wells, 25 fields per well). Dendritic marker MAP2 is in green. (D) Neuronal culture stained for GFAP (red), an astroglial marker, and TuJ1 (green). On an average of 4404±2727 cells per well, 27.6%±8.2 of them are positive for GFAP (n=12 wells, 25 fields per well). For all images, nuclei are stained with DAPI (blue).

**Figure S.5:**
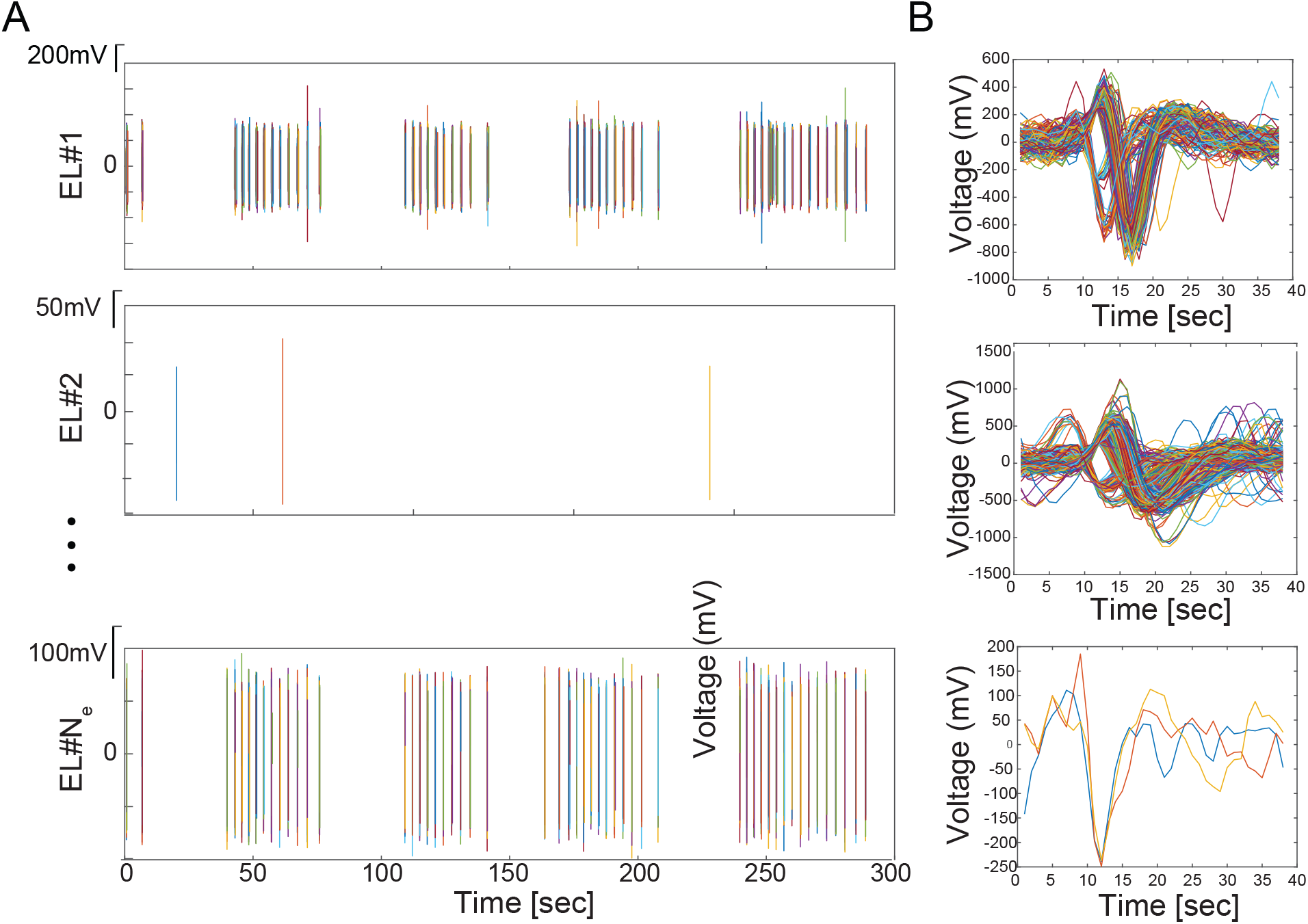
Examples of electrophysiological recordings from iPSC networks coupled to Multielectrode Arrays. (Left) Spike trains recorded by three different electrodes in one of the MEA’s well from all detected neurons. Different colors are only used to indicate unsorted spikes in the signals. (Right) Overlaid detected spike waveforms from the corresponding signal on the left side panel.

**Figure S.6:**
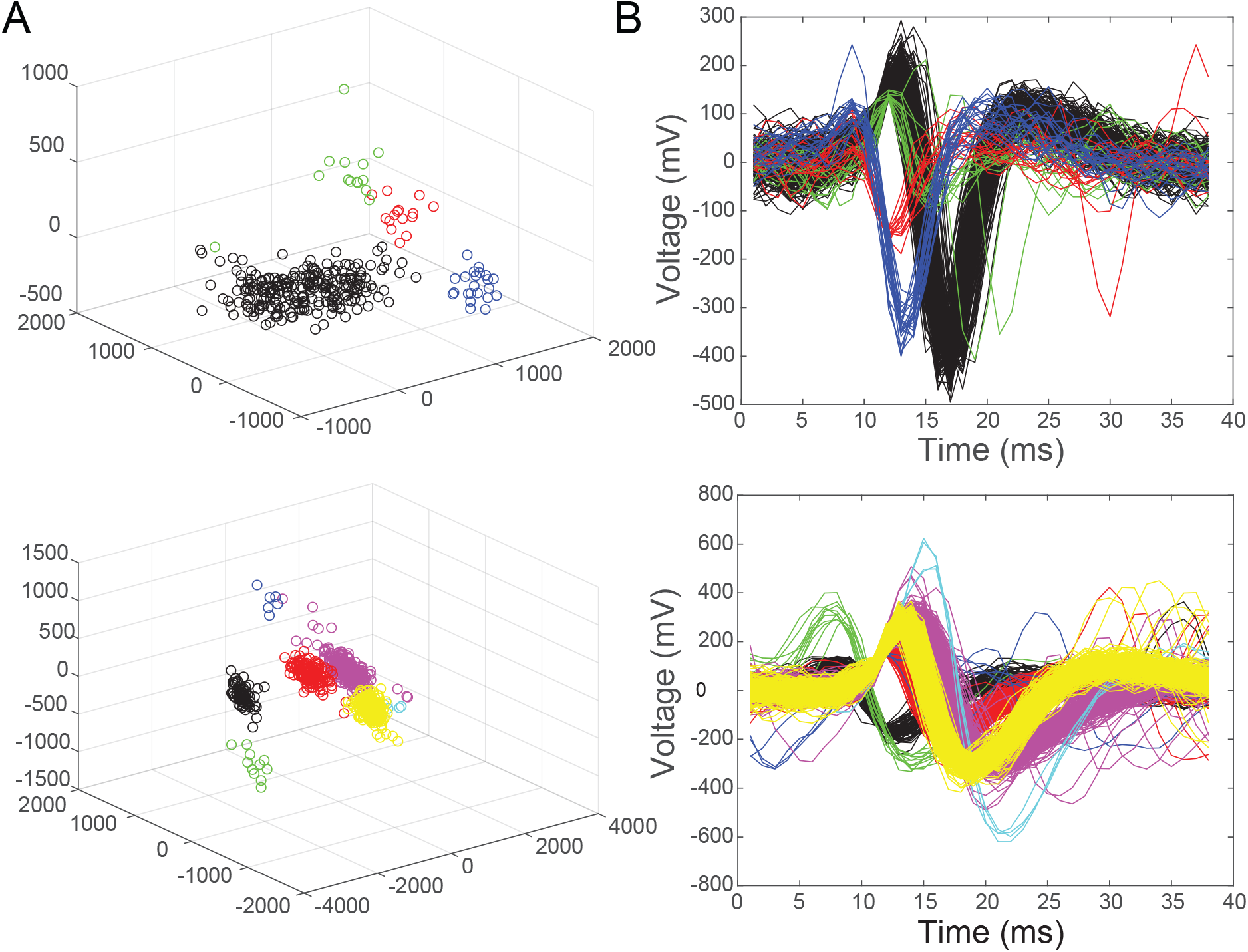
Examples of spike sorted data. Data were taken from the recordings reported in Figure S.5 (top and middle row). PCA was used to extract similar features in the spikes’ dataset. Spikes with similar features were grouped into clusters via a *k*-means clustering approach. This consisted of partitioning *n* observations into *k* clusters in which each observation belongs to the cluster with the nearest mean. (Left) The detected clusters after PCA analysis and *k*-means clustering. Different colors stand for different clusters of spike signals, each cluster belonging to a different neuron. (Right) The corresponding sorted spikes. Similar waveforms belong to the same recorded neuron.

